# Decoding the molecular interplay of endogenous CD20 and Rituximab with fast volumetric nanoscopy

**DOI:** 10.1101/2023.10.09.561472

**Authors:** Arindam Ghosh, Mara Meub, Dominic A. Helmerich, Patrick Eiring, K. Martin Kortüm, Sören Doose, Markus Sauer

**Affiliations:** Department of Biotechnology and Biophysics, Biocenter, University of Würzburg; Würzburg, Germany; Rudolf Virchow Center, Research Center for Integrative and Translational Bioimaging, University of Würzburg; Würzburg, Germany; Department of Internal Medicine II, University Hospital Würzburg, Würzburg, Germany

## Abstract

Elucidating the interaction between membrane proteins and antibodies requires fast whole-cell imaging at high spatiotemporal resolution. Lattice light-sheet (LLS) microscopy offers fast volumetric imaging but suffers from limited spatial resolution. DNA-PAINT achieves molecular resolution but is practically restricted to two-dimensional imaging due to long acquisition times. Here, we introduce two-dye imager (TDI) probes, manifesting negligible background and amplified fluorescence signal upon transient binding, enabling ∼15-fold faster imaging. Using a combination of TDI-DNA-PAINT and LLS microscopy on B cells, we reveal the oligomeric states and interaction of endogenous CD20 with the therapeutic monoclonal antibody rituximab (RTX), unperturbed by surface effects. Our results demonstrate that B cells become polarized, and microvilli stabilized by RTX binding. These findings, we believe, will aid rational design of improved immunotherapies targeting tumor-associated antigens.

## Introduction

Fast three-dimensional (3D) imaging of whole cells with high spatial resolution presents challenges to existing optical imaging techniques. Selective plane illumination microscopy such as lattice light-sheet (LLS) microscopy allows researchers to generate multi-dimensional images of samples with minimal photodamage at high speeds, thus enabling the visualization of dynamic processes in living specimens at diffraction-limited resolution (*1,2*). On the other hand, DNA-based point accumulation for imaging in nanoscale topography (DNA-PAINT) achieves molecular resolution but can be applied only to fixed samples and requires long imaging times because of the background signal originating from freely diffusing imager probes in solution (*3,4*). Several key benefits of DNA-PAINT include easy multiplexing via sequential imaging of targets of interest with different imager strands and resilience to photobleaching of fluorophores as they are constantly replenished by freely diffusing imager strands in solution (*5*). Furthermore, DNA-PAINT enables accurate sub-10 nm fluorescence imaging because it avoids energy transfer interactions between fluorophores separated by less than 10 nm (*6*). Recently, 3D volumetric imaging of cells with a spatial resolution of up to ∼60-70 nm in lateral and ∼100-110 nm in axial direction has been demonstrated by combining LLS microscopy and PAINT (*2,7*). However, a major issue of PAINT remains to be the strong background signal, limiting probe concentrations to 0.1-0.5 nM. Low concentrations, however, result in long durations of data acquisition, typically in the range of a few hours per imaging plane and up to several days for densely labeled whole cells imaged by LLS microscopy (*7*).

Lately, efforts have been made to engineer energy transfer based fluorogenic imager probes to improve DNA-PAINT imaging (*8,9*). Here, we exploit probes end-labeled with two identical dyes that form non-fluorescent dimers in the unbound state, for faster DNA-PAINT imaging (*10*). We tested multiple organic dyes and identified ATTO Oxa14 as a superior-performing fluorophore for the design of self-quenched DNA probes that efficiently form non-fluorescent H-dimers (Fig. 1A and figs. S1-S2) (*11,12*). The underlying concept involves the formation of quenched imager probes that show minimal fluorescence in the unbound state and complete recovery of the fluorescence signal upon transient binding to the complementary docking strand (Fig. 1B). We term this modality as two-dye imager DNA-PAINT (TDI-DNA-PAINT).

**Fig. 1.**
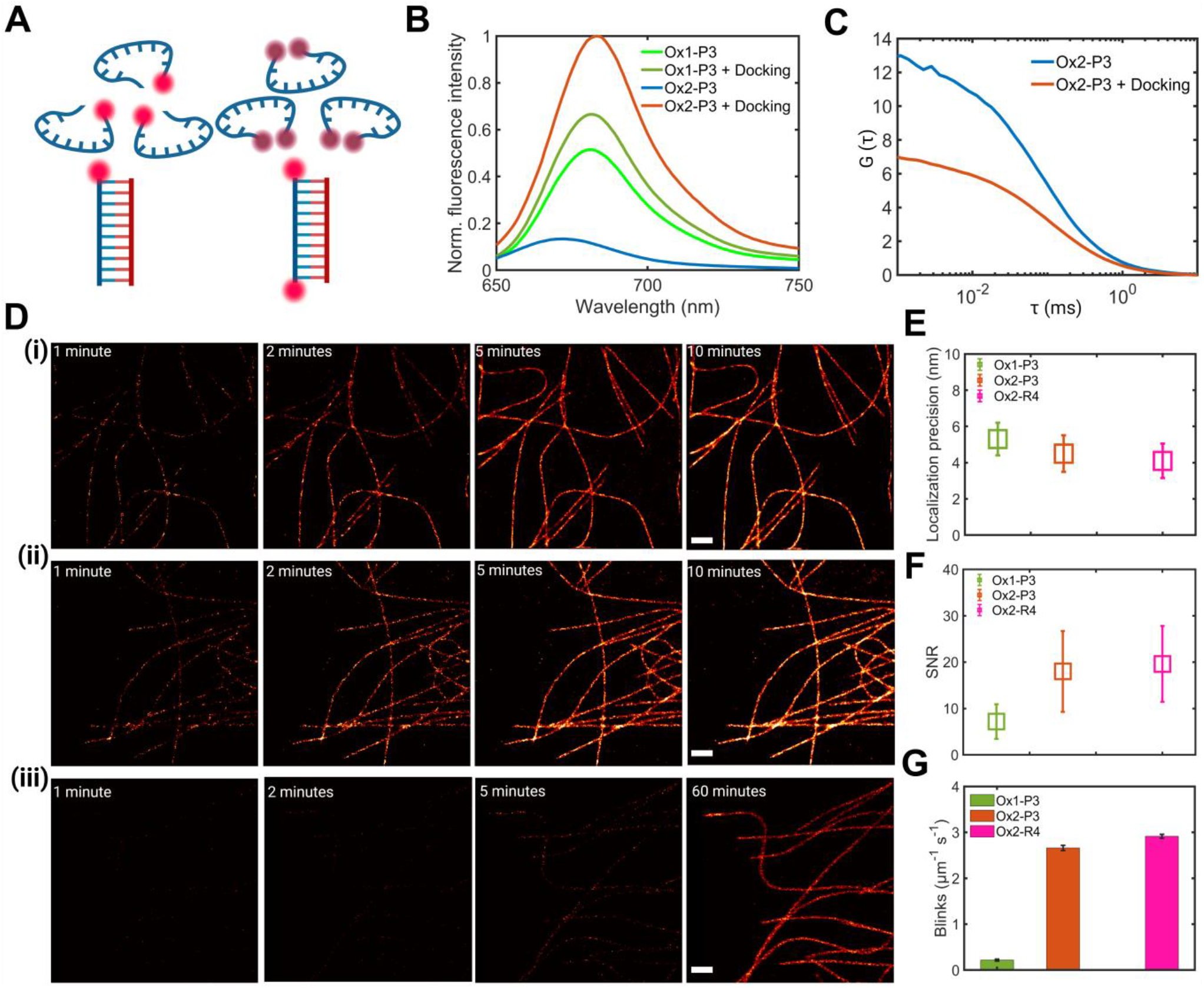
2D TDI-DNA-PAINT imaging. (**A**) Classical DNA-PAINT with single-labeled imager strands (left) and TDI-DNA-PAINT workflow with dual-labeled Ox2 imager strands (right). (**B**) Fluorescence spectra of Ox1-P3 and Ox2-P3 in unbound state and fluorescence dequenching after adding an excess of docking strands. (**C**) FCS measurement of Ox2-P3 before (blue) and after (red) adding an excess of docking strands. (**D**) TDI-DNA-PAINT imaging of microtubules using (**i**) 5 nM of Ox2-P3 strands and (ii) 5 nM of Ox2-R4 strands, as well as (iii) 100 pM of Ox1-P3 strands for direct comparison to classical DNA-PAINT. Reconstructions shown for 1, 2, 5, and 10 min. (i and ii) and for 1, 2, 5, and 60 minutes of image acquisition (iii). Scale bars, 1 μm. (**E**) Localization precisions (nm) after 5 min of TDI-DNA-PAINT imaging (Ox2-P3: red and Ox2-R4: magenta) were 4.5 ± 1.0 nm (mean ± std) and 4.1 ± 0.9 nm, respectively, whereas classical DNA-PAINT (Ox1-P3: green) experiment yielded a precision of 5.3 ± 0.9 nm after 60 minutes of data acquisition. (**F**) Comparison of the signal-to-noise ratio (SNR) from experiments shown in (**D**). (**G**) Number of blinking events per μm per second for Ox1-P3 (green), Ox2-P3 (red), and Ox2-R4 (magenta) probes.

## Results

### Faster imaging with TDI-DNA-PAINT

For the first demonstration of TDI-DNA-PAINT, we selected the routinely used classical DNA-PAINT imager strand P3 (10 nucleotides) (*3*) and the speed optimized R4 (7 nt) (*4*) and labeled them with one or two ATTO Oxa14 dyes. In the remainder of the manuscript, we will refer to the dual- and single-labeled imagers as Ox2-(P3 or R4) and Ox1-(P3 or R4), respectively. Ensemble fluorescence emission spectra of Ox2-P3 in the absence and presence of a 10^4^-molar excess of docking strand showed an 8.7-fold increase in fluorescence intensity upon binding (Fig. 1B). Ox2-R4 probes displayed a similar fluorescence quenching efficiency (fig. S3A). Furthermore, we confirmed the formation of non-fluorescent H-dimers between the two terminal ATTO Oxa14 dyes by fluorescence correlation spectroscopy (FCS). Here, we assumed that the conformational flexibility of the imager strand permits contact formation of the two dyes. The halving of the FCS amplitude upon addition of an excess of docking strands demonstrates that on average ∼ 50% of all dual-labeled probes form non-fluorescent H-dimers with a lifetime that is longer than the diffusion time through the confocal detection volume (∼1 ms) (Fig. 1C). In addition, shorter-lived dimer formation (indicated by the correlation decay on μs time scales) and collisional quenching reduces the fluorescence of undocked imager strands.

Efficient fluorescence quenching of ATTO Oxa14 dual-labeled probes in the unbound state allowed us to use Ox2-P3 and Ox2-R4 at substantially higher concentrations (fig. S3B) for imaging microtubules in COS7 cells. We could use Ox2-P3 at a concentration of 5 nM, whereas classical DNA-PAINT with Ox1-P3 had to be conducted at 100 pM concentrations (movies S1-S12). Five minutes of imaging resulted in 31,400, 27,100, and 2,490 detected localizations for the two TDI-DNA-PAINT imager strands Ox2-R4 and Ox2-P3, and the classical DNA-PAINT imager strand Ox1-P3, respectively, for an image area covered with comparable filament density (Fig. 1D). For classical DNA-PAINT, the mean localization precision from three independent 60 min measurements with Ox1-P3 was found to be 5.3 ± 0.09 nm (mean ± std). In contrast, after 5 min of TDI-DNA-PAINT imaging, we obtained localization precisions of 4.4 ± 0.1 nm and 4.1 ± 0.09 nm for Ox2-P3 and Ox2-R4 imagers, respectively (Fig. 1E) *(13)*. This corresponds to an improvement factor of 1.20 and 1.29 for Ox2-P3 and Ox2-R4 probes, respectively. The higher localization precision results from the higher signal-to-noise (SNR) of dual-labeled imager probes in the bound, unquenched state and the higher fluorescence intensity in the bound state due to the presence of two fluorophores (Fig. 1F and fig. S3C). Interestingly, the attachment of two fluorophores per imager strand has only little influence on the binding time (on-time) demonstrating that TDI-DNA-PAINT can be easily implemented using established imager sequences (fig. S3D). For a more precise estimation of the imaging speed, we computed the number of localizations (blinks) detected for TDI-DNA-PAINT and DNA-PAINT per micrometer of filament and per unit time (Fig. 1G). In case of DNA-PAINT (Ox1-P3), we detected 0.2 ± 0.02 blinks μm^-1^s^-1^ (mean ± sem), whereas for TDI-DNA PAINT at 50-fold higher imager concentration we detected 2.6 ± 0.06 and 2.9 ± 0.04 blinks μm^-1^s^-1^ for Ox2-P3 and Ox2-R4 probes, respectively, resulting in a 13-fold and 14.5-fold increase in effective imaging speed (Fig. 1G). Furthermore, we utilized the speed-optimized Ox2-R4 probe for imaging a DNA origami with four docking strands separated by 18 nm (fig. S3E-H) *(6)*. The well-defined two-dimensional DNA origami sample with 4 docking sites allowed us to use higher TDI-DNA-PAINT imager concentrations of up to 25 nM (movie S13). Already after 5 minutes imaging with Ox2-R4, we detected 24-35 localizations per DNA origami and could successfully resolve the 18 nm inter-fluorophore distance (fig. S3E-H). On average, we detected an inter-fluorophore distance of 18.9 ± 2.2 nm (mean ± sem) at an acquisition time of 5 minutes.

### Whole cell LLS-TDI-DNA-PAINT

Since TDI-DNA-PAINT provides higher spatial resolutions, lower background, higher SNR (Fig. 1F and fig. S3B), and a substantial reduction in acquisition time, we concluded that it will be the ideal method for whole-cell super-resolution fluorescence imaging by LLS microscopy. We utilized a commercially available inverted LLS microscope equipped with high-power lasers and a light sheet thickness of 1.8 μm. For 3D TDI-DNA-PAINT of whole COS7 cells immunolabeled for microtubules, we exploited a new approach for engineering the point-spread-function introducing astigmatism (Fig. 2A-C, fig. S4 and movie S14) (*14*). Using a lower TDI-DNA-PAINT imager probe concentration of 2.5 nM to accommodate for the width of astigmatic patterns, we detected 21.3 million localizations for Ox2-R4 and 17.5 million localizations for Ox2-P3 within a 4-hour acquisition time from an acquired volume of 87 μm × 72 μm × 8.8 μm (movies S15-S17). Reconstructed images of COS7 cells imaged with Ox2-R4 (Fig. 2A) and Ox2-P3 (fig. S5) imager probes demonstrate the practical feasibility of fast, 3D volumetric LLS-TDI-DNA-PAINT. For Ox2-P3, we achieved localization precisions of 24 ± 12 nm (mean ± std) along the x - axis and 59 ± 22 nm along the z - axis, whereas for Ox2-R4, the localization precisions were even slightly higher, at 14 ± 5 nm and 53 ± 15 nm along the x- and z – axis, respectively (Fig. 2D). The achieved overall localization precisions are comparable to those previously reported but are substantially lower than those achieved in our 2D experiments due to the lower numerical aperture (NA) of 1.0 in our commercial inverted LLS microscope. While individual microtubule filaments far from the coverslip could be visualized at high spatial resolution using LLS-PAINT, the localization precision was worse in the central regions of the cells due to higher emitter densities (Fig. 2C) *(7)*.

**Fig. 2.**
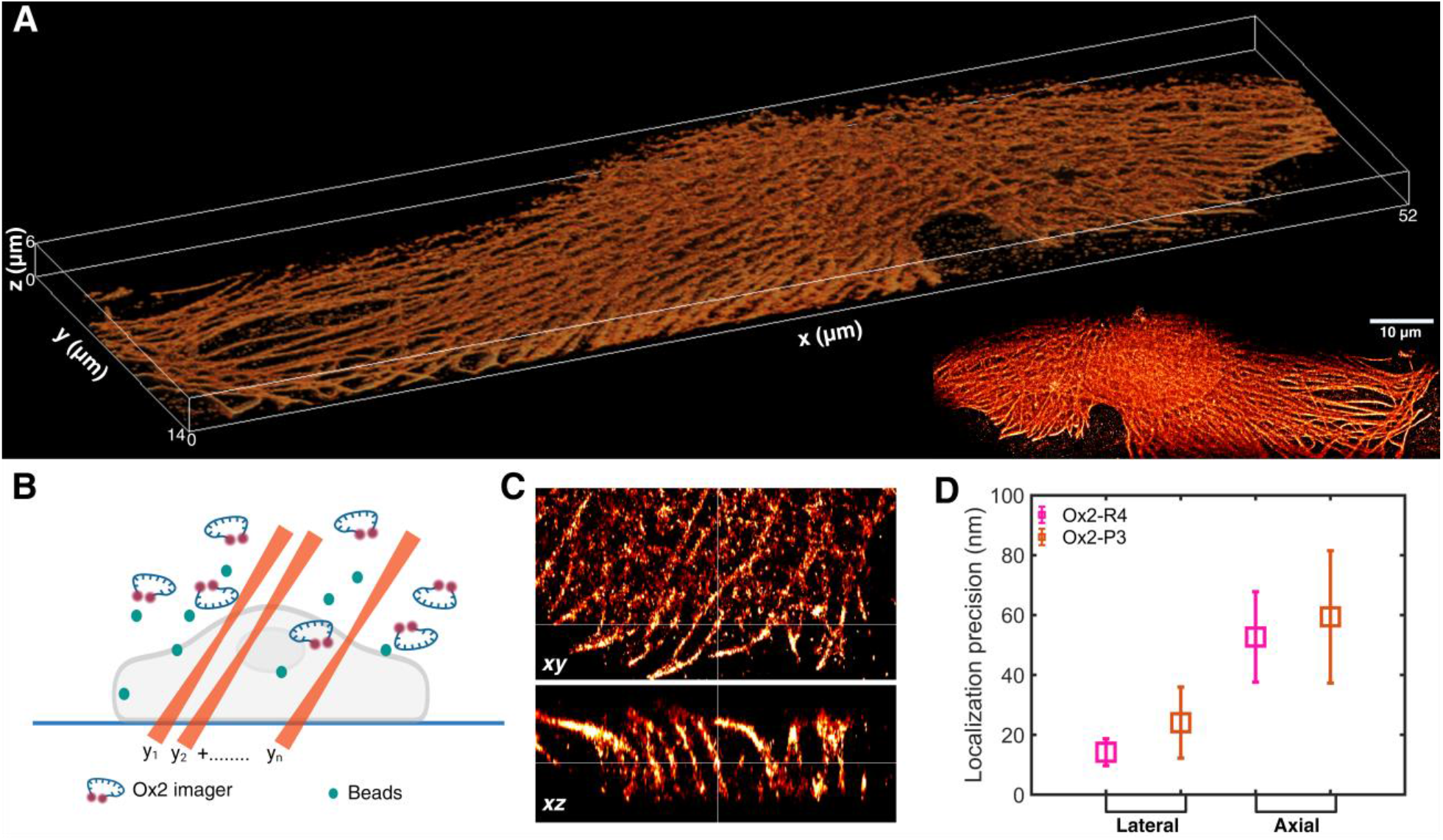
3D LLS-TDI-DNA-PAINT imaging of cellular microtubule networks. (**A**) Whole-cell volume rendering of a representative COS7 cell shows the microtubule network from TDI-DNA-PAINT imaging with the Ox2-R4 probe. The bounding box is 52 μm × 14 μm × 6 μm. The inset illustrates a maximum *z* projection of the same cell after deskewing but before rotation to the actual coverslip coordinates. (**B**) 3D LLS TDI-DNA-PAINT workflow, a cell labeled with docking strands is fixed inside a glass-bottomed well and fluorescent beads are co-incubated for PSF calibration. Ox2-P3/R4 imager probes are used at a concentration of 2.5 nM. Using a light sheet of the dimensions of 100 μm × 1.8 μm, the sample is imaged by acquiring a time series of 2,500 frames per position along the skewed *y* axis and a total of 35 positions with a step size of 500 nm (∼0.25 μm along true z of the coverslip coordinates). (**C**) Orthogonal views with *xy* - (top) and *xz* - (bottom) profiles of a representative ROI. (**D**) Localization precisions achieved from whole-cell LLS TDI-DNA-PAINT imaging for the cells shown in (**A**).

### Molecular organization of endogenous CD20 on B cells in complex with rituximab

Since LLS-TDI-DNA-PAINT enables fast 3D volumetric imaging of cells with virtually molecular resolution, we used it to investigate the organization of endogenous CD20 membrane receptors when bound to the therapeutic monoclonal antibody rituximab (RTX). The integral membrane protein CD20 is a B cell specific marker that is targeted by monoclonal antibodies (mAb) for the treatment of B cell related malignancies and autoimmune disorders. RTX was the first approved therapeutic mAb for cancer and continues to be the reference for next generation mAbs (*15-17*). Even though RTX has been in clinical use for two decades knowledge about its mechanism of action, i.e. how it activates the immune system to kill B cells remains poorly understood. The proposed lines of attack include direct cell death, Fc receptor effector functions through antibody-dependent cellular cytotoxicity (ADCC), phagocytosis and complement-dependent cytotoxicity (CDC) (*18*). However, there is no clear understanding of how the molecular interaction of RTX with CD20 (or its successor mAbs) triggers B cell depletion. In addition, knowledge about the oligomeric states of CD20 in the plasma membrane remains elusive. Available evidence suggests that CD20 associates into homo-oligomers of various stoichiometries and complexes with other proteins (*19,20*). One mode of action proposes that RTX activates CDC by oligomerization of the mAb’s Fc region to increase the binding avidity of complement component 1q (C1q), which is a hexa-headed molecule that optimally binds hexameric Fc arrangements (*21,22*). Indeed, recent studies performed with CD20 in detergents and in CHO cells transiently transfected with mEGFP-CD20 indicated the formation of CD20 dimers that are concatenated by RTX to linear, chain-like or hexameric, ring-like structures that might recruit C1q (*4,23,24*).

The fact that RTX does not merely concatenate CD20 but more generally rearranges other intracellular and membrane-bound B cell proteins, suggests that rearrangement is required for depletion of the cell by the immune system (*25*). We therefore investigated mAb-induced rearrangements of endogenous CD20 on the plasma membrane of Raji B cells by TDI-DNA-PAINT. Therefore, monoclonal antibodies labeled with docking strands were used to image the basal membrane of adherent cells in contact with the coverslip using TDI-DNA-PAINT (movies S18-S21). To depict the distribution of CD20 in the plasma membrane of Raji B cells we performed labeling experiments with 5-10 μg/ml RTX and 2H7 at 4°C on ice and room temperature as well as with various incubation times (2-30 min). 2H7 binds to the almost identical epitope of CD20 but at lower binding affinity (*18,26*). The obtained images show CD20 monomers and oligomers on the B cell membrane indicating that CD20 is natively present in oligomers or RTX induces weak concatenation of CD20 even at 4°C (Fig. 3A and fig. S6). TDI-DNA-PAINT images of the weaker binding 2H7 show similar characteristics albeit at significantly lower extent (Fig. 3A and fig. S7). Incubation of B cells with RTX at room temperature results in stronger crosslinking of CD20 and the appearance of filaments and higher order oligomers on the basal membrane of B cells after not more than 2 min of incubation time (fig. S6). In contrast, treatment with 2H7 shows a significantly lower concatenation efficiency (Fig. 3A and fig. S7).

**Fig. 3.**
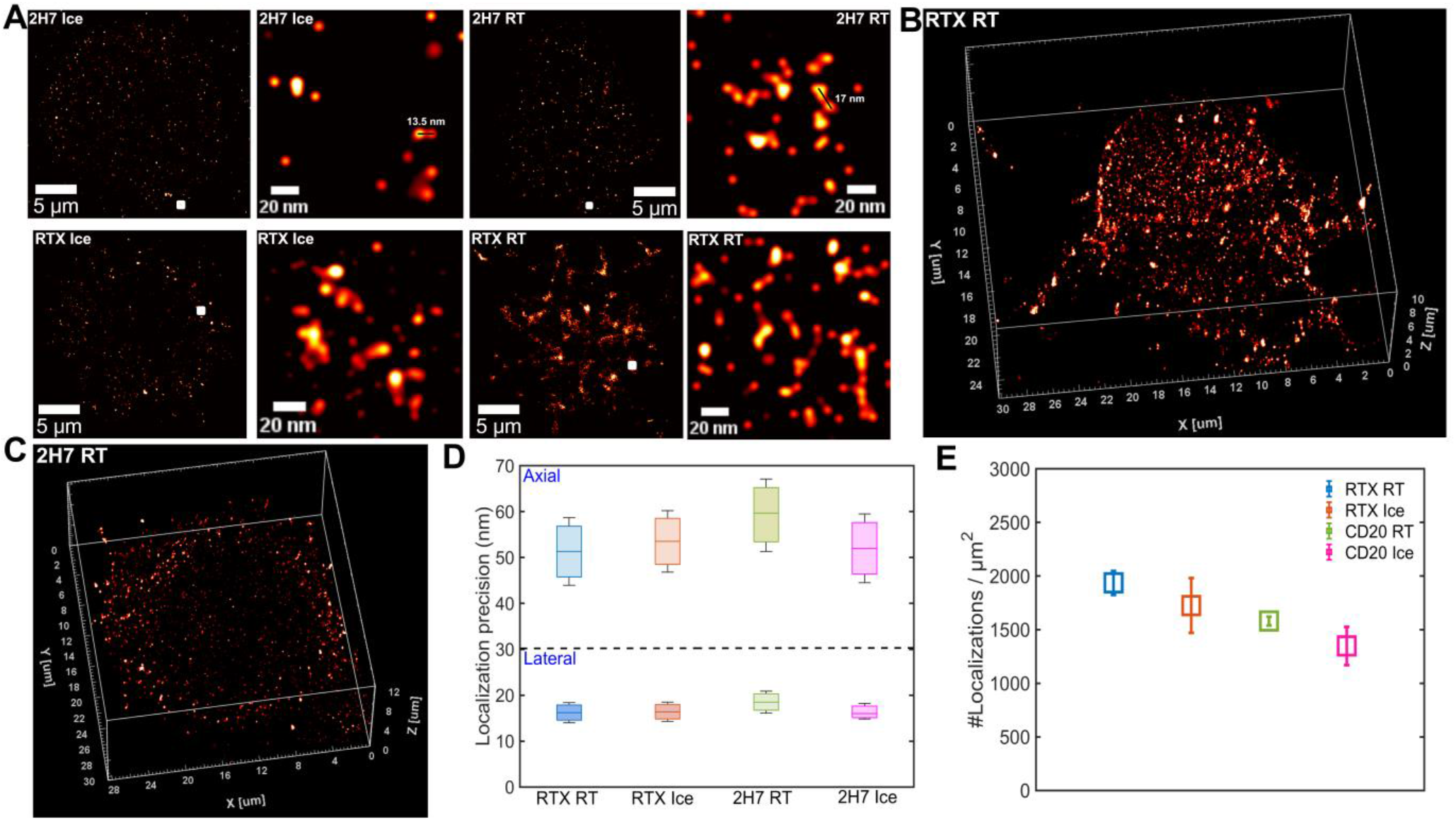
2D and 3D LLS-TDI-DNA-PAINT imaging of CD20 in Raji cells. (**A**) 2D TDI-DNA-PAINT reconstructions of basal membrane of Raji cells depict the nanoscale organization of CD20 labeled with RTX/2H7 at RT and on ice. Imaging was performed with 5nM Ox2-R4 imagers. Representative ROIs for each condition show monomers, possibly dimers (two examples shown), and oligomers in case of 2H7, whereas larger, filamentous, and higher-order oligomers can be seen for RTX. (**B**) 3D LLS TDI-DNA-PAINT volume rendering of CD20/RTX in a typical Raji cell labeled at RT. Concatenated CD20 in long membrane protrusions are observed. The bounding box is 30 μm × 25 μm × 10 μm. (**C**) same for CD20/2H7 labeled Raji cell at RT. The bounding box is 28 μm × 30 μm × 12 μm. Unlike in **(B)**, distributions of CD20 monomers, dimers, and oligomers, with a few short protrusions can be seen. Examples of cells labeled on ice are provided in fig. S8. (**D**) Axial and lateral localization precisions achieved from whole-cell LLS TDI-DNA-PAINT imaging of Raji cells under various labeling conditions are shown. **(E)** Average number of localizations per μm^2^ of a cell under various conditions are plotted.

Since CD20 is known to strongly accumulate on finger-like membrane protrusions (microvilli) of B cells with a diameter of a few hundred nanometers and a length of up to 4 μm (*27,28*), the appearance of linear, chain-like structures on the basal membrane of cells could be explained by the existence of microvilli on insufficiently adhered cells (*29*). Since the degree of adherence of a suspension cell on a coverslip varies considerably, imaging of the basal membrane is extremely error prone. This results in a strong heterogeneity observed for the distribution of CD20/RTX signals on basal B cell membranes in contact with the coverslip (Fig. 3A and fig. S6). Therefore, we performed 3D volumetric LLS-TDI-DNA-PAINT imaging with RTX and 2H7 to visualize both, the basal and apical membrane of Raji B cells (Figs. 3B-C, fig. S8, movies S22-S25). Notably, in these experiments we achieved localization precisions of <20 nm and <60 nm in lateral and axial directions, respectively (Fig. 3D). The generated 3D images clearly demonstrate the accumulation of CD20/RTX signals on microvilli, validating our hypothesis that the filamentous structures observed in 2D TDI-DNA-PAINT experiments are an artifact caused by B cell microvilli getting sandwiched between the basal membrane and the coverslip. Interestingly, the CD20/2H7 signals appear to be overall lower with 1580 ± 81 (mean ± sem) and 1348 ± 178 detected localizations per μm^2^ compared to 1963 ± 112 and 1725 ± 256 detected localizations per μm^2^ for CD20/RTX stained at RT and on ice, respectively, and the length of B cell microvilli substantially shorter (Figs. 3B, C, 3E and fig. S8).

### RTX polarizes B cells and stabilizes microvilli

Microvilli are finger-like membrane protrusions of the actin cytoskeleton that play an important role in antigen surveillance and sensing of B cells (*28*). To fully understand the interaction of therapeutic antibodies with plasma membrane receptors that trigger B cell depletion, real-time 3D imaging with high spatiotemporal resolution is required. Therefore, we investigated the dynamic interaction of CD20 with different concentrations of dye labeled RTX and 2H7 on living Raji B cells by two-color LLS microscopy (*30,31*). To enable the visualization of the microvilli dynamics, actin was labeled with a fluorogenic live-cell stain. The obtained movies in the presence of 5 and 10 μg/ml RTX unequivocally prove the point: CD20/RTX complexes are predominantly localized on microvilli, that expand upon RTX exposure (Figs. 4A, C, and D). Simultaneously, incubation with RTX induces polarization of the cells, i.e. CD20/RTX complexes move actively to one side of the B cell (Fig. 4A, figs. S9, S10, and movies S26-S30). Time-resolved investigation of the actin signal after RTX treatment shows rapid reorganization of actin ruffles into microvilli protrusions (Fig. 4A, figs. S9, S10) (*32*) reaching sizes of up to 7.3 μm (median length 2.8 μm, Figs. 4C-D)

**Fig. 4.**
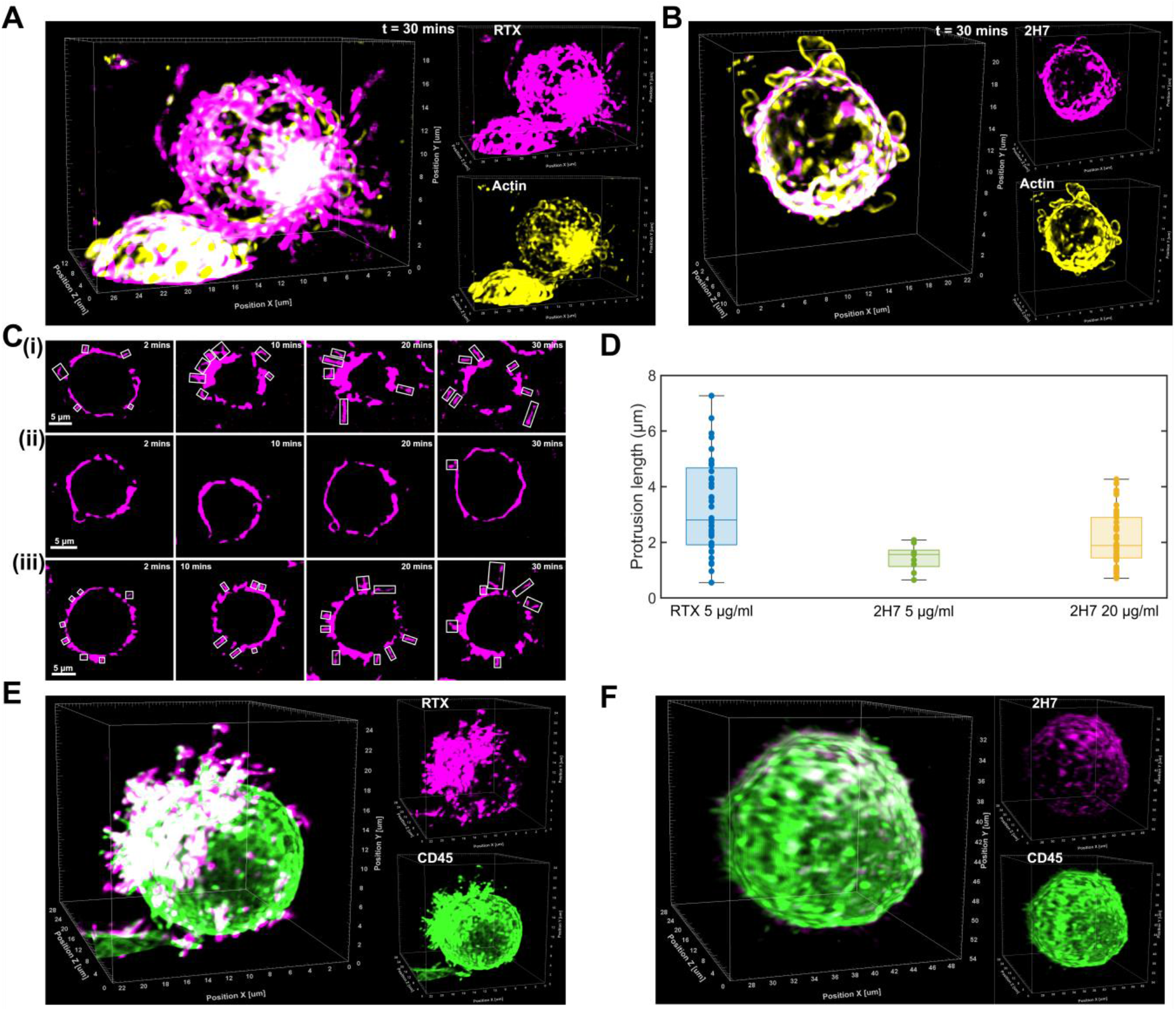
Two-color LLS imaging in Raji cells. (**A**) Live-cell LLS imaging of CD20 and actin in Raji cells with 5 μg/mL RTX-AF647 (magenta) and SPY-555 for actin (yellow). Deconvolved LLS image after 30 minutes of antibody addition manifest microvilli stabilization of varied dimensions enriched in CD20 clusters along with polarized CD20 aggregation. Individual channels are illustrated in inset (Movie S26). (**B**) Live-cell LLS imaging of CD20 and actin in Raji cells with 5 μg/mL 2H7-AF647 (magenta) and SPY-555 for actin (yellow). Image shows actin-enriched membrane ruffles continues to exist after 30 minutes of 2H7 incubation (Movie S27), in contrast to (**A**). (**C**) 2D images from an exemplary plane at different timepoints after antibody addition for the cells **(i)** after RTX addition as shown in **(A), (ii)** after 2H/ addition as shown in **(B)**, and **(iii)** after 2H7 addition (fig. S13A). Identified microvilli are demarcated with rectangles. (**D**) Protrusion length calculated for the cells in **(C)** for 46, 9, and 44 identified microvilli in case of 5 μg/mL RTX, 5 μg/mL 2H7, and 20 μg/mL 2H7 respectively. **(E)** Two-color LLS imaging with 5 μg/mL RTX-AF647 (magenta) and CD45-CF568 (green) in Raji cells. Polarized, higher order CD20 aggregates and their accumulation in microvilli are visualized. **(F)** Two-color LLS imaging with 5 μg/mL 2H7 -AF647 (magenta) and CD45-CF568 (green) in Raji cells. The B cells do not show aggregation or long microvilli enriched in CD20.

Treatment with 5 μg/ml 2H7 results in overall weaker binding and lower accumulation of CD20/2H7 complexes on microvilli accompanied by substantially shorter microvilli (median length 1.6 μm and maximum length 2.1 μm) and less pronounced polarization of B cells (Figs. 4B-D, figs. S11, S12, and movies S26-30). In contrast, elevating the concentration of 2H7 leads to longer microvilli protrusions (median length 1.9 μm and maximum length 4.3 μm) and B cell polarization, but even at high concentrations of 20 μg/ml, 2H7 does not match the magnitude of effects induced by 5 μg/ml of RTX (Figs. 4C-D and fig. S13). Two-color LLS microscopy experiments on living Raji B cells treated with dye-labeled RTX and an antibody directed against the transmembrane phosphatase CD45 confirm that RTX polarizes the cell and CD20/RTX complexes strongly accumulate on microvilli and stabilize them in an outstretched form while CD45 is excluded from microvilli tips (Fig. 4E and figs. S14-15) (*29,33*). Here again, our data corroborate that 2H7 induces much weaker effects (Fig. 4F and fig. S15).

## Discussion

To examine the molecular organization of target proteins on whole cells and understand how they interact with therapeutic antibodies to induce the depletion of malignant or autoimmune B cells, high resolution imaging methods are required that are, regrettably, not yet available. Here, by using an oxazine dye with superior H-dimer formation tendency we could successfully tackle the main challenges associated with DNA-PAINT imaging, long acquisition times and high background levels. The substantially reduced fluorescence background of ATTO Oxa14 labeled self-quenched imager probes enables the use of higher imager concentrations and consequently up to 15-fold higher imaging speed with higher localization precision as compared to classical DNA-PAINT. The reduced background fluorescence and accelerated imaging speed provides the basis for whole-cell 3D LLS-TDI-DNA-PAINT imaging within a few hours. Since TDI-DNA-PAINT works with established DNA-PAINT sequences it can be easily adopted and does not require the design of optimized docking and imager strands as necessary for FRET-based approaches *(8,9)*. Thus, TDI-DNA-PAINT can also be advantageously used to speed up multiplexed DNA-PAINT imaging approaches *(5)*. Among the screened fluorophores, TDI-DNA-PAINT with ATTO Oxa14 outperformed Cy5, ATTO 655, and tetramethyl rhodamine (fig. S2). However, a careful screening for other fluorophores with stronger H-dimer formation efficiency might pave the way for even more efficient and faster LLS-TDI-DNA-PAINT at molecular resolution.

TDI-DNA-PAINT with docking strand-labeled RTX on Raji B cells clearly showed monomeric, dimeric and multimeric CD20/RTX complexes of different geometry (Fig. 3A and fig. S6) corroborating the strong concatenation efficiency of RTX (*19-24*). 3D LLS-TDI-DNA-PAINT experiments on Raji B cells expressing endogenous CD20, however, revealed that linear, chain-like CD20/RTX structures on the basal membrane of cells may well be explained by B cell microvilli sandwiched between the basal membrane and the coverslip that are stabilized by RTX binding (Figs 3B, C and fig. S6). Since CD20 is highly expressed on microvilli of B cells (*25,26*) concatenation of CD20 by RTX is highly effective. This results in local high concentrations of Fc fragments organized in various geometries along the microvilli (*21,22*). It is very likely that these CD20/RTX oligomeric structures are required to recruit complement component 1q (C1q) (*4,23,24*).

However, our two-color live cell LLS microscopy experiments clearly indicate that the oligomerization state of CD20/RTX complexes is just one piece in a multifarious puzzle. 3D live-cell imaging clearly demonstrates that CD20/RTX complexes accumulate on microvilli that rapidly move to one side on the B cell membrane within a few minutes (Figs. 4A, E and figs. S9, S10). In addition to B cell polarization that might augment triggering of ADCC by natural killer (NK) cells (*25*) microvilli are stabilized and their protrusion length is substantially longer when compared to treatment with the weaker binding 2H7 (Figs. 4C-D). In B cells, antigen surveillance and sensing are enabled by microvilli that homogeneously cover the cell surface. The stabilization of CD20/RTX complexes on microvilli and the movement of microvilli towards one side of the B cell indicates a dramatic reorganization within the cell membrane resembling the formation of an immunological synapse (*25*). In B cells, the immunological synapse is the site for antigen recognition and downstream signaling and it might thus have important implications for ADCC (*30-32,34*). Indeed, there is evidence that macrophages and neutrophils conjugate with antibody opsonized target cells (*35*). Undoubtedly, our data show that whole-cell imaging techniques allow us to visualize the molecular interaction of CD20 and RTX unperturbed by surface effects. The combination of LLS microscopy and TDI-DNA-PAINT provides the high spatiotemporal resolution required to capture a complete picture that can guide the development of improved immunotherapies.

## ACKNOWLEDGEMENTS

The authors thank Lukas Kapitein, Ralf Jungmann, and Mike Heilemann for fruitful discussions.

## Author contributions

A.G. and M.S. conceived the project and wrote the manuscript. A.G. and M.M. performed 2D TDI-DNA-PAINT. A.G. performed 3D LLS-TDI-DNA-PAINT imaging experiments, analyzed data, and generated the figures for the main manuscript. M.M. did cell culture, prepared microtubule samples, did experiments on DNA origami, and helped in manuscript editing. D.A.H synthesized DNA origami platforms, prepared the final sample, and helped in manuscript editing. S.D. helped in data analysis and manuscript editing. M.K. and P.E. supported RTX experiments.

## Data and Materials availability

The data that support the findings of this study will be provided by the corresponding author upon reasonable request.

## SUPPLEMENTARY MATERIALS

Materials and Methods

Figures S1 to S15

Tables S1 and S2

Movies S1 to S30

References and Notes (36-40)

