## Supporting Material for "Decoding the molecular interplay of endogenous CD20 and Rituximab with fast volumetric nanoscopy"

**The PDF file includes:**

**Materials and Methods**

**Figs. S1 to S15**

**References and Notes**

**Other Supplementary Material for this manuscript includes the following:**

**Tables S1 and S2**

**Movies S1 to S30**

### Materials and Methods

#### Imager and docking DNA strands

Methyltetrazine modified docking strands P3’: MetTet-5′-TTTCTTCATTA-3′ and 7× R4’: MetTet- 5’-ACACACACACACACACACA-3’ (Table S1); imager probes P3: 5’-GTAATGAAGA-3’ single and double labeled with the dyes Cy5, ATTO655, TMR, and ATTO Oxa14 (Ox2-P3). R4: 5’-TGTGTGT-3’ double labeled with ATTO Oxa14 (Ox2-R4), and ATTO 520 were obtained commercially (biomers.net GmbH).

#### Imaging buffers

Imaging buffer for TDI-DNA-PAINT imaging of microtubules in COS7 cells and CD20 in Raji cells: PBS (Sigma-Aldrich, #D8537-500 ML) + 500 mM NaCl (Sigma-Aldrich, #S5886) at pH 7.4. Imaging buffer for DNA-Origami samples: PBS (Sigma-Aldrich, D8537-500 ML) + 5 mM TRIS (Invitrogen, #AM9855G) + 50 mM $MgCl_2$ (AppliChem, #A4425, 0500) + 1 mM EDTA (Sigma-Aldrich, #E1644-250G) + 0.05% Tween20 (ThermoFisher, #28320) adjusted to pH 7.6.

#### Cell culture

African green monkey kidney fibroblast-like cells (COS7, Cell Lines Service GmbH, Eppelheim, #605470) were cultured in DMEM (Sigma-Aldrich, #D8062) containing 10% FCS (Sigma-Aldrich, #F7524), 100 U/mL penicillin and 0.1 mg/mL streptomycin (Sigma-Aldrich, #P4333) at 37 °C and 5% $CO_2$. Cells were grown in standard T25-culture flasks (Greiner Bio-One). Human B-lymphocytes (Raji, Cell Lines Service GmbH, #300359) were cultured in RPMI-1640 (Sigma-Aldrich, #R8758) containing 10% FCS, 100 U/mL penicillin and 0.1 mg/mL streptomycin at 37°C and 5% $CO_2$. Cells were maintained at a maximum density of ~$2x10^6$ cells/mL in standard T25 culture flasks (Sarstedt # 83.3910.502).

#### Antibody modification for (LLS) TDI-DNA-PAINT imaging

For labeling of goat anti-rabbit IgG (H+L) secondary antibody (Invitrogen, #31212), anti-human CD20 (2H7, BioLegend, #302302) and Rituximab with PEG4-TCO, an excess of TCO-PEG4-NHS was used. Antibody labeling was performed at 20 °C for 4 h in labeling buffer (100 mM sodium tetraborate (Fulka, #71999), pH 9.5) following the manufacturer’s standard protocol. The antibodies were reconstituted in labeling buffer using 0.5 mL spin-desalting columns (40K MWCO, ThermoFisher, #87766). The conjugated antibodies were purified using spin-desalting columns and stored in PBS. Finally, the antibody concentration was determined by UV–vis absorption spectrometry (Jasco V-650).

**Tubulin staining for (LLS) TDI-DNA-PAINT**

COS7 cells were seeded at concentrations of $2.5 \times 10^4$ cells/well into eight-well chambered glass slides with a high-performance cover glass bottom (Cellvis, C8-1.5H-N). Cells were grown until proper adhesion at 37 °C and 5% $CO_2$. Cells were washed with pre-warmed (37°C) PBS and permeabilized for 2 min using 0.3% glutaraldehyde (GA) + 0.25% Triton X-100 (EMS, #16220 and ThermoFisher, #28314) in pre-warmed (37°C) cytoskeleton buffer (CB) which consists of 10 mM MES ((Sigma-Aldrich, #M8250), pH 6.1), 150 mM NaCl, 5 mM EGTA (Sigma-Aldrich, #03777), 5 mM glucose (Sigma-Aldrich, #G7021), and 5 mM $MgCl_2$ (AppliChem, #A4425, 0500). Following permeabilization, cells were fixed with a pre-warmed (37 °C) solution of 2% GA for 10 min. After GA- induced fixation, cells were incubated for 5 min in 100 mM glycine (Ajinomoto, #G5417) to stop fixation followed by washing twice with PBS. Samples were reduced with 0.1% sodium borohydride (Sigma-Aldrich, #71320) in PBS for 7 min to avoid background fluorescence, and again washed three times with PBS. After blocking with 5% BSA (Roth, #3737.3) for 30 min, cells were incubated with primary antibody (rabbit anti-alpha tubulin antibody, Abcam, #ab18251, 5 μg/mL) for 1 h at 20°C followed by three washing steps with PBS. Next, cells were incubated with secondary goat anti-rabbit IgG labeled with PEG4-TCO (10 μg/mL) for 1 h at 20 °C and again washed three times with PBS. After washing, cells were incubated with MetTet-5′-TTTCTTCATTA-3′ (P3') or MetTet- 5'-ACACACACACACACACACA-3' (7× R4') single-stranded docking DNA strands (biomers.net GmbH, custom-made, 10 μg/mL) for 15 min at RT. The Methyltetrazine modification allowed the attachment of docking strands to secondary antibody via 'click chemistry'. As a last step, cells were washed three more times with PBS (Sigma-Aldrich, #D8537-500 ML) and stored at 4 °C until imaging.

**Rituximab (RTX) and 2H7 staining of Raji cells for (LLS) TDI-DNA-PAINT**

Raji cells were seeded at concentrations of $2.5 \times 10^5$ cells/well into poly-D-lysine (PDL) (Sigma-Aldrich, #P6407) coated eight-well chambered glass slides with a high-performance cover glass bottom ((8-well Chambered Coverglass System #1.5 High Performance Cover Glass (0.17 ± 0.005 μm), Cellvis) and allowed to adhere in the incubator. After the cells had successfully attached to the glass surface, they were stored at room temperature for 5 min and were transferred on ice for further 5 min in case of incubation on ice. The media was replaced with PBS containing 5 µg/mL TCO-labeled anti-CD20 2H7 (2H7-TCO, Biolegend, clone 2H7, #302302) or RTX (RTX-TCO, RTX was kindly provided by the Pharmacy of the University Hospital of Würzburg) antibody and incubated for 30 min if not indicated otherwise. After washing, cells were incubated with the docking strand (7×R4': MetTet- 5'-ACACACACACACACACACA-3') for 30 min and again washed with PBS. For fixation, cells were incubated for 15 min with 4 % methanol-free formaldehyde (FA) (ThermoFisher, #28906) and 0.2 % glutaraldehyde (GA) (Sigma-Aldrich, #G5882-10×1ML), again washed with PBS and stored at 4°C in the dark or directly measured.

**Antibody-dye conjugation for LLS imaging of CD20 and CD45**

RTX and anti-CD20 2H7 (Biolegend, clone 2H7, #302302) were self-labeled with AF647-NHS (Thermofisher, #A20006) and CD45 (Biolegend, clone HI30, #304002) with CF568-NHS (Merck, #SCJ4600027). The antibodies were labeled at RT for 2 h in 100 mM sodium bicarbonate (Fisher Scientific, 144-55-8, pH 8.5) following manufacturers standard protocol. Briefly, 50 µg of antibody was reconstituted in sodium bicarbonate buffer using 0.5 mL spin desalting columns (40K MWCO, Thermofisher, #87766). To get an average DOL of 2-3 a 5× dye excess was used. All antibodies were purified and washed using additional spin desalting columns to remove unbound dye. Finally, antibody concentration was determined measuring absorption (A) at 280 nm and 568 or 647 nm with a nanophotometer respectively and calculated according to the following formula with ε being the extinction coefficient and CF the correction factor of the dye at 280 nm.

$$\text{Protein concentration (M)} = \frac{A_{280} - (A_{dye}\ x\ CF)}{\varepsilon^{protein}}\ x\ dilution\ factor$$

$$\text{Degree of labeling (DOL)} = \frac{A_{max}}{\varepsilon^{dye}\ x\ protein\ concentration\ (M)}\ x\ dilution\ factor$$

**Immunostaining of CD20 and CD45**

Raji cells were seeded on 0.1 mg/mL PDL (Sigma-Aldrich, #P6407) coated Cellvis chamber slides (8-well Chambered Coverglass System #1.5 High Performance Cover Glass (0.17±0.005 µm), Cellvis) and incubated with 5 µg/mL of either anti-CD20 RTX-AF647 or 2H7-AF647 and anti-CD45 HI30-CF568 on ice for 45 min. Cells were washed once with ice cold PBS prior fixation with 4% methanol-free formaldehyde / 0.2% glutaraldehyde for 15 min (Thermofisher, 28906 & Sigma-Aldrich, G5882) and stored at 4 °C until imaging.

### Live-cell actin and CD20 staining in Raji cells

For two-color live-cell LLS imaging of actin and CD20 in Raji cells, actin was stained with a bright, orange, fluorogenic, and non-toxic F-actin stain SPY555-FastAct™ (Spirochrome AG, #SC205). Actin labeling was performed following the manufacturer's protocol: At first, 1000× stock solution is prepared by adding 50 µL of anhydrous DMSO to the SPY555-FastAct™ vial. The stock was prepared using newly opened and anhydrous DMSO. After use, this solution was stored at -20 °C. 1000× stock solution of SPY555-FastAct™ was diluted to 1× in pre-warmed cell culture medium (RPMI-1640 (Sigma-Aldrich, #R8758) containing 10% FCS, 100 U/mL penicillin and 0.1 mg/mL streptomycin). Raji cells were seeded on PDL-coated 8-well chambers (Coverglass System #1.5 High Performance Cover Glass (0.17 ± 0.005 µm), Cellvis) and allowed to adhere for 30 min at 37 °C and 5% $CO_2$ in the incubator. After 30 min, cells were allowed to equilibrate at room temperature (RT) for 15 min, after which the culture medium was replaced by 1× staining solution freshly prepared ensuring that all the cells are covered with the solution. The cells were placed in the incubator (37 °C in a humidified atmosphere containing 5% $CO_2$) for 1 h. After the labelling, cells were not washed since the probe is fluorogenic and directly transferred to the microscope for imaging. RTX or 2H7 was added at desired concentration (5/10/20 µg/mL) to respective wells at the microscope stage and measurement was started immediately after addition of RTX/2H7 to Raji cells.

### DNA origami surface preparation

DNA-origami single-molecule surfaces were prepared in eight-well chambers (following the exact described protocol *(6)* containing high performance cover glass (Cellvis, C8-1.5H-N). The wells were washed once with PBS (Sigma-Aldrich, #D8537-500ML) before treatment with 2% Hellmanex (Hellma, 9-307-011-4-507) for 1 h. Following the treatment, the chambers were washed thrice with PBS. Next, the surfaces were incubated with 1 M KOH (Fulka, #06005) for 20 min followed and washed three times with PBS. Next, the surfaces were incubated with 10% polyethylene glycol 400 (Fulka, #81170) overnight at 4 °C. On the following day, the surfaces were rinsed thrice with PBS and the chambers were incubated with 0.5 g/l BSA-Biotin (ThermoFisher, #29130) in PBS overnight at 4 °C. Next day, the chambers were washed again thrice with PBS before incubation with 0.5 g/l Neutravidin (ThermoFisher, #31050) in PBS for 20 min. The surfaces were washed three times with PBS and incubated with purified DNA-origami solution, 1:5 diluted in

PBS + 50 mM $MgCl_2$ (AppliChem, #A4425,0500) for 10 min. The prepared samples were washed at least thrice with PBS + 50 mM $MgCl_2$ before imaging.

**Preparation of DNA-origami structures**

Design, hybridization and quality control of DNA-origami rectangular structures were carried out as described earlier *(6)*, using the same DNA sequences to hybridize the final 18 nm spaced structure. Briefly, sequences were designed with caDNAno v.2.2.0 and stability calculations of the origami designed were performed using CanDO *(36, 37, 38)*. We obtained the trans-cyclooctene (TCO) modified staple strands (biomers.net GmbH) commercially, biotinylated strands (Sigma-Aldrich), and unmodified staple strands (Merck KGaA). The scaffold DNA used in this study was phage M13mp18 derivate DNA type p7560 (tilibit nanosystems, M1-32). We performed the hybridization reaction by mixing 10 nM scaffold DNA with 15× surplus of unmodified staple strands and 30× surplus of modified staple strands in hybridization buffer (5 mM tris (hydroxymethyl) aminomethane (TRIS) (Merck, 1.08382.2500), 5 mM sodium chloride (NaCl) (Sigma-Aldrich, S5880-1KG), 1 mM ethylene diamine tetra-acetic acid (EDTA) (Sigma-Aldrich, #E1644-250G) and 12 mM magnesium chloride ($MgCl_2$) (AppliChem, #A4425,0500)) using a ThermoCycler (C1000 Thermal Cycler, BioRad) by applying a linear thermal gradient of $-1\ °C\ min^{-1}$ from 90 to 4 °C. In case of origami samples for TDI-DNA-PAINT, TCO modified staple strands were used. After hybridization, the origami nanostructures were incubated with a 10× surplus of docking strands (P3': MetTet-5′-TTTCTTCATTA-3′ or 7xR4': MetTet- 5'-ACACACACACACACACACA-3') per incorporated TCO-staple for click reaction overnight at 4 °C. Finally, the clicked and hybridized samples were purified by gel electrophoresis using a 1.5% agarose gel (Sigma-Aldrich, #A9539-500G) in 1× TBE buffer, consisting of 4.5 mM TRIS, 4.5 mM Boric acid, 10 mM EDTA. A 0.5× TBE was used as running buffer. A final concentration of 12 mM $MgCl_2$ was added to all TBE buffers. A few µl of hybridized and clicked origami structures were used as references and mixed with an intercalating dye. The rest of the hybridized origami samples were not mixed with the intercalating dye. All samples were incubated with loading dye (10 mM TRIS (Merck, 1.08382.2500), 60% glycerol (v/v) (Merck, 1.37028.1000) and 0.03% bromophenol blue (w/v) (Carl Roth, T116.1). Electrophoresis was done at 70 V, using a programmable d.c. voltage source (PowerPac Basic, BioRad), for ~ 2 h in water/ice bath. Finally, the agarose gel was cut into two pieces to separate the reference band and the samples not incubated with intercalating dye. The reference band was marked at an ultraviolet transilluminator (UST20M-8E, INTAS). Afterwards the marked gel was combined with the not illuminated part of the gel and not illuminated DNA origamis were cutted out accordingly. Extracted agarose gel parts were cutted several times and purified via Freeze N′ Squeeze columns (Freeze N′ Squeeze, 7326165, BioRad) according to the manufacturer's instructions using a benchtop centrifuge (Biofuge fresco, Heraeus) at 13,000*g*. For all measurements, the DNA origami was prepared freshly and imaged immediately.

**Ensemble absorbance and fluorescence spectroscopy**

UV / visible absorbance spectra and steady-state fluorescence emission spectra of fluorescently labeled imager probes were recorded using a Jasco V650 absorbance spectrometer and Jasco FP-6500 spectrofluorometer respectively. For absorbance spectra, 500 nM Ox2 and Ox1 imagers were used (fig. S1). In case of fluorescence spectra, 1 nM imager strands and respective docking strands were used in $10^4$ molar excess (10 µM). Fluorescence emission spectra of P3 labeled with 2×ATTO655, 2×Cy5, and 2×TMR, 2×Ox2-R4, 2×ATTO 520 R4 imager strands (fig. S2). Samples were measured in a 3 mm path-length cuvette (Hellma, 105.251-QS) in PBS (Sigma-Aldrich, #D8537-500ML) containing 500 mM NaCl adjusted at pH 7.4.

**Fluorescence correlation spectroscopy (FCS)**

FCS measurements on Ox2-P3 probes (Fig. 1C) were carried out using a commercial time-resolved confocal fluorescence microscope setup (MicroTime200, PicoQuant) as described previously *(39)*. Imager strands were used at a concentration of 1 nM dissolved in PBS (Sigma-Aldrich, #D8537-500ML) containing 500 mM NaCl adjusted at pH 7.4, while the complementary docking strand was $10^4$ times higher i.e., 10 µM.

**TDI-DNA-PAINT imaging**

TDI-DNA-PAINT imaging of microtubules in COS7 cells (Fig. 1D), DNA origami structures (fig. S3E-H), and CD20 in Raji cells (Fig. 3A, fig. S6-S7) were performed utilizing an inverted wide-field fluorescence microscope (IX-71; Olympus). For excitation of Ox2 probes, a 640 nm diode laser (Cube 640-100C, Coherent) in combination with a clean-up filter (laser clean-up filter 640/10, Chroma) was used. The laser beam was focused onto the back focal plane of the oil-immersion objective (100×, NA 1.51; Olympus). Emission light was separated from the illumination light using a dichroic mirror (HC 560/659; Semrock) and spectrally filtered by a band-pass filter (FF01-679/41-25, Semrock).

Images were recorded with an electron-multiplying CCD camera chip (iXon DU-897; Andor). Pixel size for data analysis was measured to be 104 nm. For classical DNA-PAINT and TDI-DNA-PAINT experiments on cellular microtubule networks, 5 nM Ox2-P3 / Ox2-R4 probe and 100 pM Ox1-P3 probe was added to the respective glass-bottomed chamber wells containing COS7 cells fixed and immunolabeled with corresponding MetTet-P3' docking strand (Fig. 1D). For measurements on Raji cells on CD20 labeled via RTX or 2H7, 5 nM Ox2-R4 probe was used. Blinking movies (Supplementary movies S1-S12, S18-S21) were recorded for representative 128 × 128-pixel areas (for microtubule in COS7 cells) and 256 × 256-pixel areas or similar (for CD20 with RTX or 2H7 in Raji cells) with TIRF excitation with an exposure time of 100 ms (frame rate 9.96 Hz) and irradiation intensity of 0.8 kW/cm$^2$. Images were recorded for 18k frames (30 min) for each dataset (Table S2). We recorded images of DNA-origami samples using a different setup but with same optical parameters and components as for the previous one. We used this second setup as it is a more robust "drift-free" system in our laboratory and can be beneficial for origami samples. We added 25 nM of Ox2-R4 probes to the well containing DNA-origami nanostructures labeled with MetTet -7×R4' docking strand incubated in the imaging buffer for origami. Images were recorded for 128 × 128-pixel areas (Movie S3, Table S2) with exposure times of 30 ms for 90k frames at 1.1 kW/cm$^2$. Pixel size for data analysis was measured to be 128 nm.

**Data analysis for 2D TDI-DNA-PAINT**

TDI-DNA-PAINT data for microtubules and DNA-origami were analyzed using the MATLAB-based Super-resolution Microscopy Analysis Platform (SMAP) *(40)*. Using SMAP, data analysis was done by following the steps: loading of camera images, filtering, background estimation, finding of candidate molecules, fitting, and saving of the results. First, we loaded the raw blinking movies into the SMAP GUI, which was followed by manually setting up the camera parameters. Next, under the *Localize\Peak Finder* tab, we set the parameters for the initial estimation of single molecule positions, such as filtering cutoff with intensity values. To ensure selecting the brightest localizations, we select a higher cutoff value. After previewing an exemplary frame, we chose the Gaussian 2D fitting tab *PSF free* for localizing the full image set. The analysis returns coordinates such as x, y and z, number of photons per localization, background values, localization precisions, frame in which localization was found etc. Grouping (merging) of localizations persistent over several frames is performed after fitting. We can manually set the following grouping parameters dX (maximum distance two localizations can be apart) and dT (maximum number of dark frames between localizations). We grouped the localizations by setting dT = 1, and limiting the localization precision range from 0-15 nm, which filters out localizations with worse precision. On applying these grouping parameters, SMAP calculates the maximum distance from the localization precision of the two candidate localizations. That means that localizations are only grouped if it is likely they stem from the same event. We also quantified the background signal from different concentrations of Ox2-P3/R4 probes from 100 pM to 10 nM, and 5 nM appeared to be the most optimal concentration considering it had very similar background as compared to 100 pM Ox1-P3 (fig. S3B). We also tested if the attachment of two fluorophores to the imager P3 introduced any change to binding time (on-time). Figure S3D depict the on-time comparison between Ox1-P3, Ox2-P3, and Ox2-R4. Figures S3C and S3E-S3H presents photon yields, localization precision, background signal, and detected origami structures from TDI-DNA-PAINT experiments with Ox2-R4. The reported distances from origami experiments were from $N$ = 33 values. For CD20 TDI-DNA-PAINT datasets, analysis was performed following the same workflow as described and reconstructed images from 10 min acquisition is illustrated in Fig. 3A and figs. S6-S7.

**Whole-cell lattice light sheet (LLS) TDI-DNA-PAINT imaging**

Here, we present a detailed description of microscopy setup, sample preparation, imaging, and data analysis for the datasets presented in this study. The following workflow mostly recapitulates the description from Iwanski et al. *(14)*.

**Microscopy and data analysis setup**

Zeiss lattice light-sheet 7 equipped with

a. 13.3x/ 0.4 N.A. excitation lens, 44.83x /1.0 N.A. detection lens
b. Emission notch filter 420-470, 503-546, 576-617, 656-750 nm
c. Camera: ORCA-Fusion canal sCMOS (Hamamtsu Photonics), final pixel size 145 nm
d. Lattice light-sheet 100 x 1800 (length [µm] x thickness [nm])
e. 488, 561, 640 nm excitation laser lines

For our measurements, we used only 640 nm laser with the power density of 3.3 kW / cm$^2$ (as specified by the manufacturer) at the specified light sheet.

i) ZEN software for the acquisition control and image deskewing
ii) ZEN macro for automating acquisition.

iii) FIJI (Fiji is just ImageJ v1.53q) equipped with the following plug-ins:

a. DoM (Detection of Molecules v.1.2.4, https://github.com/ekatrukha/DoM_Utrecht) software for detection and image reconstruction

b. FTM2 (Faster Temporal Median filter, https://github.com/HohlbeinLab/FTM2) used to filter out rapid moving (diffusing) imagers

c. Register NDFFT (Registration of ND images using Fast Fourier Transform, v0.10.5, https://github.com/ekatrukha/RegisterNDFFT) software for averaging of images used here for finding an average PSF for z-calibration

d. Correlescence (v.0.0.4) (https://github.com/ekatrukha/Correlescence) software for spatio-temporal correlation analysis used here for slice-to-slice drift correction.

iv) Image processing FIJI macros for batch single molecule detection, PSF extraction and averaging, super-resolution image reconstruction and drift correction. These run the above plug-ins with pre-defined settings for each slice.

v) Final localization tables analysis and reconstruction scripts (in MATLAB or Python).

vi) All plugins, macros, scripts (together with example data) used in this protocol are available from *(42)* and at https://doi.org/10.6084/m9.figshare.c.6244869

**Sample preparation**

African green monkey kidney fibroblast-like cells (COS7, Cell Lines Service GmbH, Eppelheim, #605470) immunolabeled with docking strands P3' and 7×R4' for tubulin were prepared as described before. Human B-lymphocytes (Raji, Cell Lines Service GmbH, #300359) was labeled for CD20 with 5 μg/mL of RTX/2H7 'click-labeled' with 7×R4' docking strand as described before. Additionally, for PSF calibration purpose, we added TetraSpeck beads (0.1 μm; Life Technologies #T7279) to the wells at dilution of 1:1000. The beads were sonicated prior to use. Exemplary PSF calibration is presented in Supplementary fig. S4.

**Imaging**

After focusing on the sample, we chose the "Sinc3 100x1800" light sheet followed by adjusting imaging parameters such as correction for the X and Y tilt of the coverslip, "Focus Sheet" and "Focus Waist". To introduce astigmatism, we adjusted the "Aberration control" parameter in increments of 5 μm while scanning the stage around a position of a few calibration beads. The core idea is to visually find a value that introduces astigmatic features to the PSF. At this value, the image of the PSF would transition from "horizontally elongated" to "vertically elongated" as we scroll the stage through a bead. For #1.5 high precision coverslips of 170 μm, we found the best parameter value to be 165 μm. The example of such an astigmatic PSF is shown in fig. S4 and Supplementary Movie S14. Next, we find a representative cell for imaging and scan the stage around the cell to ensure the volume around the cell has at least 5-6 isolated beads that can be later used for PSF *z*-calibration. Following this step, ROI was cropped such that it includes the whole volume of the cell with some margins around. We then set the exposure time to 100 ms and a laser power of 100%. Before imaging the selected cell, PSF z-calibration scan of beads was performed as:

a. We unchecked the "Time Series" checkbox and acquired a volumetric "Sample scan" with the stage displacement in *y* set to 0.2 μm intervals/step.
b. For this acquisition, we recommend extending the start and end scan positions of the stage beyond (~3-5 μm) the boundaries of the selected cell to include more beads.
c. From this point on, all imaging/microscope parameters were kept constant for the following time series acquisitions. We saved the acquired volume (PSF z-calibration) in a separate folder.

Next, we unselected the "Sample Scan" checkbox in the acquisition settings and selected "Time Series" with the "Number of frames per slice" as 2500 (#frames). We parked the stage at the boundary of the cell corresponding to the minimum value of the stage *y* coordinate, since the following ZEN macro acquisitions of the cell volume will move the stage to increase *y*. Using the provided '3D LLS' ZEN macro in the Macro Editor, we adjusted the following parameters:

a. *StageStepSize* This represents the stage displacement step between consecutive slice acquisitions (in μm). We used 0.8 μm that allows sufficient overlap of the PSF in *z* between adjacent slices.
b. *StageNumberOfSteps* This parameter represents the total number of slices which can be calculated by dividing the total *y* stage range covering the cell by the *stage step size*. We used 35 steps for acquisition on COS7 cells and 60 steps for Raji cells respectively.
c. *SaveFolderFile* This is a string with the full path to the folder where the data will be stored plus an initial filename template.

After registering these parameters, we started data acquisition. In the actual skewed dimension, a volume of 87 µm × 72 µm × 17.6 µm which, in coverslip coordinates translate to 87 µm × 72 µm × 8.8 µm, was acquired for the cell presented in Fig. 2A. Exemplary blinking movies are provided (Movies S15, S22-23) corresponding to the cells in Figs. 2A,

3B-C. During the acquisition, the calculated and true *y* positions of the stage appear in “Messages” window of the macro editor. We saved this log information after completion of data acquisition as a text or .csv file. This file will be required later for registration of sample drift between slices and deskewing operations.

**Data analysis**

i) First, the PSF *z*- calibration data was deskewed in the ZEN software using “Deskew” function keeping the “Processing method” equal to “Deskew”, but not “Coverslip transformation”.

ii) Extracting individual images of PSF / beads:

a. The deskewed z-calibration file was opened in FIJI to create a maximum intensity ‘Z project’.

b. We selected 3-4 bright and consistent beads that are relatively isolated by drawing a custom ROI.

c. With the ROI active, the DoM plugin was run (*Analyze->DoM ->Detect Molecules*). Using the “Preview detection” option, the intensity threshold parameter was adjusted such that only selected beads are detected within the ROI. The ‘Analysis’ was run and the plugin generates a Results Table with coordinates of the beads.

d. Next, we select the deskewed calibration z-stack and run the provided “extract_psf_batch.ijm” ImageJ macro. This will generate the extracted z-stacks of individual beads (estimated PSFs) that would be used for averaging at the next step.

iii) Calculate averaged PSF:

a. After extracting individual PSFs, *Plugins->ARegisterNDFFT->Iterative Averaging* command was executed in FIJI with the following parameters:

b. Input images: “Specify images in a folder”.

c. Initial template: “Average (center)”

d. Number of iterations: 10

e. Select “Exclude zero values”

f. Maximum shift fraction: 0.3

g. During processing, the plugin generates an average z-stack of input images and then iteratively register each individual stack to it and repeat the process.

h. Upon completion, the plugin will produce a Results table with average cross-correlation values and optimal displacements of individual stacks, together with a new 32-bit z-stack dataset containing the final averaged image of the PSF.

i. The final stack was cropped in all dimensions to remove blank z-slices and/or side artifacts.
j. The cropped stack was converted to 16-bit while carefully rescaling the intensity range.

iv) Create an astigmatism Z-calibration file:

a. We opened the averaged PSF stack in FIJI and executed *Analyze->DoM->Detect molecules* with the “Mark detected particles in overlay” option selected. It is required to scroll through the z-stack and if necessary, re-run the detection with adjusted parameters so that the PSF position is detected in as many of the z-slices as possible. The final generated Results table was saved.
b. Next, we run the *Analyze->DoM->Z axis (astigmatism)-> Make Z calibration* command with the following parameters:
c. “Make z calibration from”: Particle table (the plugin will read current Results Table from the previous step)
d. Spacing between z-planes: 100 nm. Since we acquired the calibration stack with a stage *y* displacement of 0.2 µm (200 nm), it translates to a 100 nm z slice distance in the deskewed dataset.
e. Make sure that the “Account for wobbling in X and Y” checkbox is selected and press OK.
f. The “Fit Z calibration” dialog should appear.
g. The calibration fitting of the Z range was restricted using range Zmin and Zmax parameters based on the shape of the curve displayed on the bottom left plot. Usually, we select an interval where the curve is monotonously decreasing and exclude outliers at the sides that deviate from this trend. It is possible to change the degree of the polynomial fitted to the calibration curve, but usually a third-degree polynomial is used. After clicking the “Perform fit” button, the dialog shows or updates the fitted curves (fig. S3). Once parameters are adjusted and the fits display reasonable agreement with the measured data, it is important to write down the final values of Zmin and Zmax being used. After that, by clicking “Store calibration”, the plugin saves the calibration to the FIJI/ImageJ registry (which remains there even after you restart ImageJ until a new calibration is stored).
h. To save the calibration, the *Analyze-> DoM-> Z axis (astigmatism)- > Save Z calibration* command was executed and saves as a .txt file. These calibration files can be loaded using the “Load Z calibration” command.

v) Post-z calibration, we performed batch processing of the acquired dataset using the provided "SMLM_batch_process_per_slice.ijm" ImageJ macro. This macro will process the frames acquired in each *z* slice by performing detection, drift correction, and create super-resolved reconstructions with localization files containing 3D coordinates, localization precisions, etc.

vi) The macro has the following analysis steps with corresponding parameters:

a. Single molecule detection and fitting is done with the DoM plugin. At this stage, two main parameters are considered: "PSF standard deviation" (in pixels) called *nSDPSF* in the macro and an intensity threshold expressed as the signal-to-noise ratio (SNR) of a detected particle (nSNR). For datasets with astigmatism, we find that a PSF SD value of ~1.8-2.0 and a SNR of ~4-8 work well. In general, these settings usually depend on the background level, fluorophore brightness, and wavelength used for imaging. They can be optimized beforehand by running the DoM plugin separately on a time series of a single slice and comparing the output detection quality for different settings.

b. After localizing single molecules, the macro automatically performs a calculation of their Z-coordinates using the Z-calibration currently stored in the ImageJ registry.

c. For drift correction within the time series, the DoM plugin splits the dataset into batches of nDCBatch frames (default value is 400), makes individual super-resolution reconstructions images with the pixel size of *nDCPixelSize* (in nm) and register all batch images to account for the drift.

d. After detection and drift correction, the macro generates a super-resolution reconstruction for each time series/stage position with the pixel size determined by the parameter *nPxRecon* (in nm, usually 30). Additionally, it is also necessary to specify the threshold of localization precision of particles included in the reconstruction, nMaxLocalization (in nm, default 100). The detected localizations are grouped with the parameters, i) a detected molecule off for one frame was grouped as one. ii) with a cutoff localization precision of <100 nm, the maximum allowed link distance was deduced.

e. After adjustment of the parameters, the "SMLM_batch_process_per_slice.ijm" macro was run on the folder containing the time series files for each stage position. The macro generates and saves the following indexed output: a) Results tables containing single molecule localization results, saved as a csv file with "Z_Tr_Results_" as a prefix b) Super-resolution reconstruction .tif images with "Z_Tr_Reconstruction_" as a prefix.

vii) The reconstructions produced for each slice are still in the "skewed" frame of the acquisition. So, we need to deskew the acquired volume. In order to do so, the

"Z_Tr_Reconstruction_" image series was loaded into FIJI as one stack using the *File->Import- >Image Sequence* command.

viii) Next, using the *File->Import->Results* command, the "stage_pos.csv" file produced initially by the ZEN acquisition macro is loaded. This file contains a column with the header "stage_true" that contains the proper stage *y* positions in µm. Now, we execute the "Deskew_reconstructions_stage_pos.ijm" macro which generates a new *z*-stack where super-resolution reconstructions will be deskewed. It is saved as an intermediate result.

ix) Next, we correct the sample's drift between slices (between each time series acquisition). After loading the deskewed stack from the previous step into FIJI, we run *Plugins-> Correlescence->2D cross-correlation* with the following options: a) Calculate 2D cross-correlation between: Consecutive images b) Interval between images: 1 frame c) Calculation method: FFT d) Select "Correct drift" e) Select "Limit max displacement" and fill in the maximum expected drift values between two consecutive frames in x and y (we recommend a starting value of 30 px for each).

x) This plugin uses cross-correlation and registers individual images in the stack and updates the stack. If the final result is satisfactory, the produced Results Table as "Results_DriftCorr_Correlescence.csv", else the plugin is re-run for a better drift correction. Supplementary movies S16-17, S24-25 show deskewed and drift corrected *z* stacks of the cells from Fig. 2A, 3B-C, and fig. S5.

xi) In the next step, we execute the MATLAB script "assemble_all_slice_data" script in order to combine the localization csv files from each slice into one large, updated Results table. This script will deskew coordinates and apply drift correction. In order to execute this script, we need the .csv tables with localizations from each slice, the.csv file containing the stage positions output by the ZEN macro ("stage_pos.csv"), drift correction results between slices ("Results_DriftCorr_Correlescence.csv"), and the pixel size of the reconstructions used for this. The minimum and maximum values of PSF z-calibration range (Zmin and Zmax) is also used as inputs. Finally, the script outputs a combined .csv file containing all the detections from each slice in the format of DoM plugin Results table.

xii) The provided script contains a boolean parameter *rotate* which specifies whether the coordinates of molecules will be rotated to the system of coordinates where the coverslip represents the *xy* plane (coverslip transformation) and tilt due to coverslip will be also corrected. The final angle of rotation is 30 degrees with correct setup alignment. This angle can be specified using the parameter *anglerad*. In case using the angle as 30 degrees still do not perform a correct rotation, one can also estimate the angle value following the steps described here *(42)*.

xiii) The final joint Results.csv file contain the 3D localization coordinates in the columns "X_(nm)", "Y_(nm)", "Z_(nm)" with corresponding localization precision columns having the suffix "_loc_error".

xiv) In order to create a 3D z-stack (with discrete slices) of localization data suitable for volumetric rendering algorithms, the data can be loaded to ImageJ/FIJI using *Analyze-> DoM-> Load large Results Table*. Using the *Analyze-> DoM ->Reconstruct Image* command, a set of options for the final reconstruction are available. "Pixel size of reconstructed image" specifies the *xy* pixel size, which is usually set to the mode (or median) of *xy* localization error. For a *z*-stack reconstruction, the checkbox "3D-reconstruction" is selected with the "Render as:" option equal to "Z-stack". The value of "Z-distance between slices" equal to the mode (or median) of the axial localization error. Poorly localized detections can be further filtered using the "Cut-off for localization precision" parameter (recommended value of 100 nm).

**Live-cell two-color LLS imaging of actin and CD20**

Dual-color LLS imaging of Raji cells was carried out using the Zeiss lattice light-sheet 7. Live-cell incubation with 6% $CO_2$ and 37 ˚C was maintained using the inbuilt stage top Incubator (ibidi, #12722). Actin was stained with the live-cell F-actin dye SPY555-FastAct™ (Spirochrome AG, #SC205) as described previously. RTX (5 µg/mL (Fig. 4A and fig. S9) or 10 µg/mL (fig. S10)) or 2H7 (5 µg/mL (Fig. 4B and fig. S11) or 10 µg/mL (fig. S12) or 20 µg/mL (fig. S13)) was added to the cells in the respective sample chamber right before the start of experiment. Fluorophores were excited with "Sinc3 30 × 1000" light sheets for wavelengths ($\lambda_{exc}$) 561 nm (SPY555-FastAct™) and 640 nm (RTX-AF647/2H7-AF647) via 13.3×/ 0.4 N.A. excitation objective lens. After focusing on the sample, imaging parameters such as correction for the X and Y tilt of the coverslip, "Focus Sheet" and "Focus Waist" were adjusted. The FOV in focus was scanned at 0.2 µm *y* stage position displacement in skewed dimension for a total range of 100 µm. Experiment was started right after adding RTX/2H7 to the chamber containing Raji cells and continued for 40 min with a sample scan repeating at 1 min interval. Emission light was collected through 44.83× /1.0 N.A. detection objective lens, followed by splitting to two cameras (ORCA-Fusion canal sCMOS (Hamamtsu Photonics), final pixel size 145 nm) with the aid of a 640 long-pass filter. Additionally, emission band-pass filters (576-617 and 656-750 nm) were also used before the respective cameras. Image analysis was performed using ZEN 3.9 blue software using the analysis method "Lattice light sheet": with this protocol, acquired datasets were first deconvolved with a software in-built 'Constrained Iterative' deconvolution algorithm (6 iterations), followed by deskewing the image datasets, and finally rotated to coverslip coordinates. Dual color images for RTX or 2H7 (magenta) and actin (yellow) from these measurements are shown in figs. S9-S13 with the same intensity gray value scaling for the respective channel across images for a final comparison of fluorescence signals between RTX and 2H7 and actin in different conditions tested. Movies S26-S30 depict Raji cell dynamics for 30 min at different RTX and 2H7 concentrations.

**Two-color LLS imaging of CD45 and CD20**

Raji cells were labeled with 5 µg/mL of either anti-CD20 RTX-AF647 or 2H7-AF647 and anti-CD45 HI30-CF568 and fixed after RTX/2H7 incubation following the protocol described earlier. Two-color LLS imaging was performed using the commercial Zeiss lattice light-sheet 7 setup with the same settings as described in the previous section. Dual color Raji cell volumes (CD20 (magenta) and CD45 (green)) shown in Fig. 4E-4F and figs. S14-15 demonstrate RTX accumulation in a polarized manner and its localization in membrane protrusions, corroborating our findings from live-cell LLS and LLS-TDI-DNA-PAINT measurements.

**Supplementary Table S1.** Docking strands used

| **Docking strand** | **Sequence [5' →3']** | **Modification** |
|---|---|---|
| P3 | TTTCTTCATTA | 5'-Methyltetrazine |
| R4 | ACACACACACACACACACA | 5'-Methyltetrazine |

**Supplementary Table S2.** Concentrations of imager probes, laser power, and camera settings used for spectroscopy and TDI-DNA-PAINT imaging measurements presented in this work.

| Figure | Imager | Sequence 5' →3' | Modification | Conc. | Laser Power | Exposure Time | Frames |
|---|---|---|---|---|---|---|---|
| 1D | Ox1-P3 | GTA ATG AAG A | 5'-ATTO Oxa14 | 100 pM | 0.8 kW $cm^{-2}$ | 100 ms | 18.000 |
| | Ox2-P3 | GTA ATG AAG A | 5'/3'-ATTO Oxa14 | 5 nM | 0.8 kW $cm^{-2}$ | 100 ms | 18.000 |
| | Ox2-R4 | TGT GTG T | 5'/3'-ATTO Oxa14 | 5 nM | 0.8 kW $cm^{-2}$ | 100 ms | 18.000 |
| 2A | Ox2-R4 | TGT GTG T | 5'/3'-ATTO Oxa14 | 2.5 nM | 3.3 kW $cm^{-2}$ | 100 ms | 2,500 $position^{-1}$ |
| 3A | Ox2-R4 | TGT GTG T | 5'/3'-ATTO Oxa14 | 5 nM | 0.8 kW cm-2 | 100 ms | 18.000 |
| 3B, 3C | Ox2-R4 | TGT GTG T | 5'/3'-ATTO Oxa14 | 2.5 nM | 3.3 kW $cm^{-2}$ | 100 ms | 2,500 $position^{-1}$ |
| S2E, S2H | Ox2-R4 | TGT GTG T | 5'/3'-ATTO Oxa14 | 25 nM | 1.1 kW $cm^{-2}$ | 30 ms | 90.000 |
| S4 | Ox2-P3 | GTA ATG AAG A | 5'/3'-ATTO Oxa14 | 2.5 nM | 3.3 kW $cm^{-2}$ | 150 ms | 2,500 $position^{-1}$ |

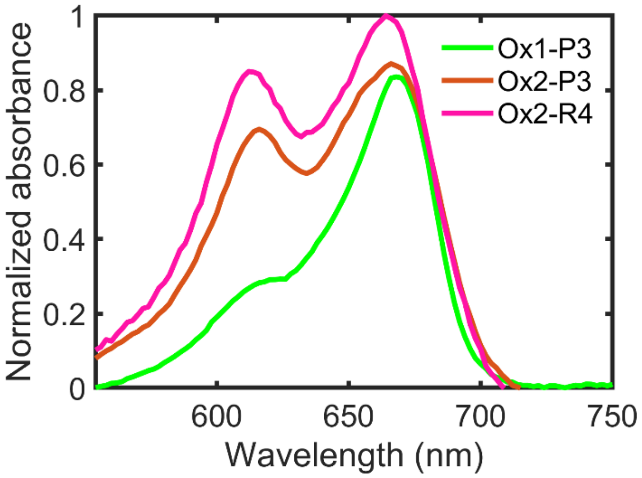


**Supplementary Fig. S1.** Normalized absorbance spectra of Ox1-P3 (green), Ox2-P3 (orange), and Ox2-R4 (magenta) imager strands recorded in PBS, containing 500 mM NaCl, pH 7.4. For Ox2-P3 and Ox2-R4 samples, a non-fluorescent H-dimer formation can be identified by the appearance of a hypsochromic-shifted shoulder in the absorption spectra, which is missing in case of Ox1-P3.

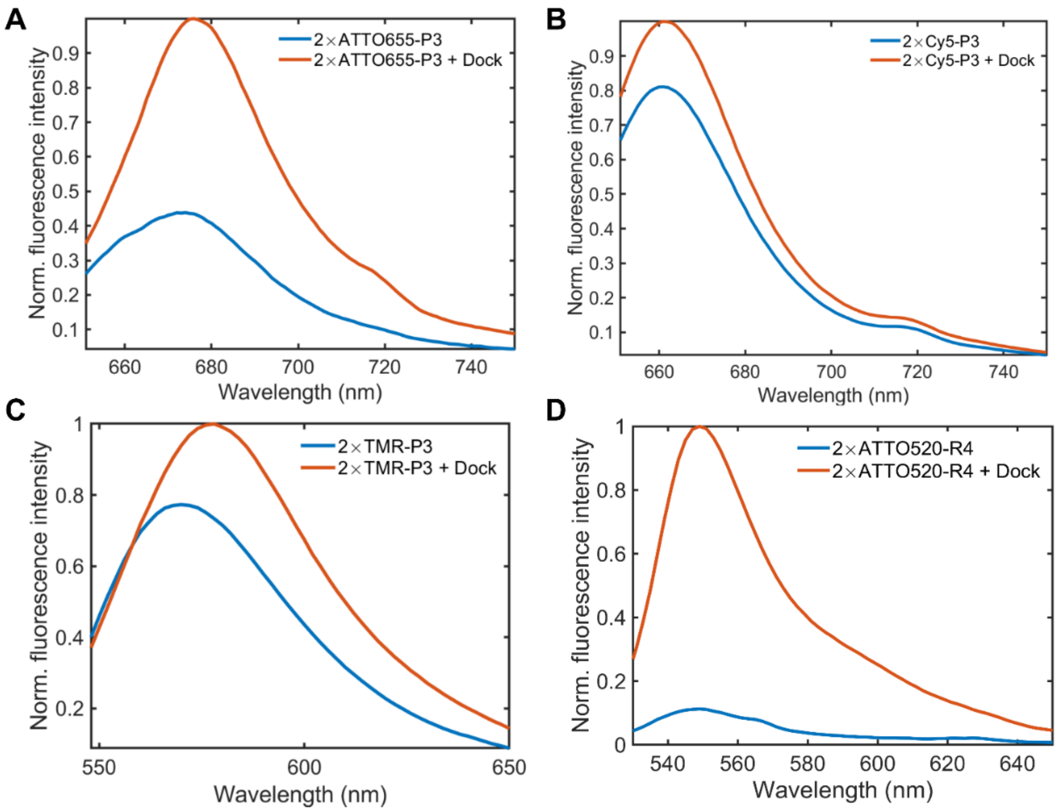


**Supplementary Fig. S2.** Normalized fluorescence emission spectra of TDI-DNA-PAINT imager probes in the absence and presence of a $10^4$-molar excess of docking strands measured in PBS containing 500 mM NaCl, pH 7.4. For the double-labeled samples (A) ATTO655-P3, (B) Cy5-P3, (C) TMR-P3 and (D) ATTO520-R4 a ~2.5-fold, ~1.25-fold, ~1.36-fold and ~9.8-fold increase in fluorescence intensity was observed, respectively.

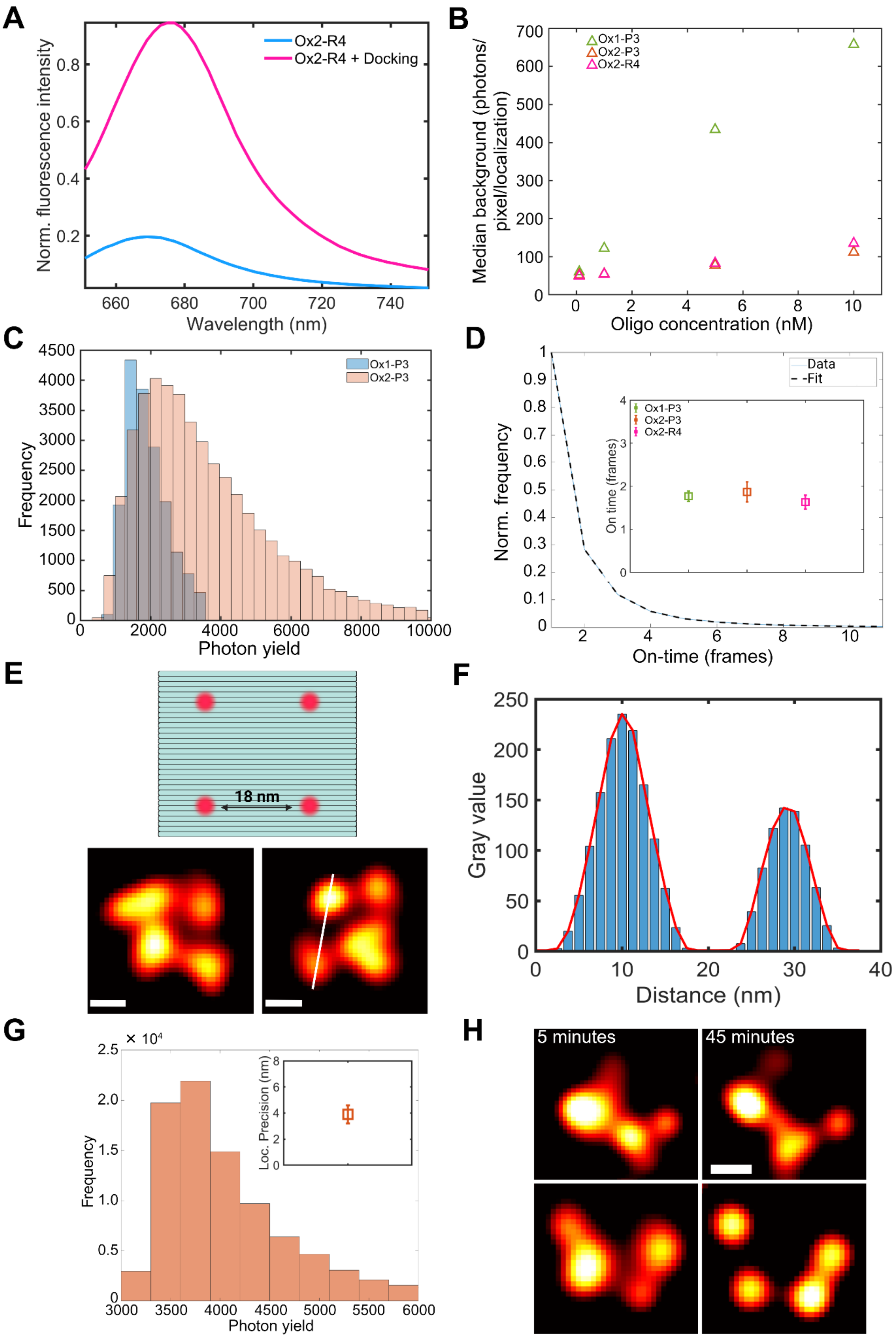
A
Ox2-R4
Ox2-R4 + Docking
Norm. fluorescence intensity
Wavelength (nm)
B
Ox1-P3
Ox2-P3
Ox2-R4
Median background (photons/ pixel/localization)
Oligo concentration (nM)
C
Ox1-P3
Ox2-P3
Frequency
Photon yield
D
Data
Fit
Ox1-P3
Ox2-P3
Ox2-R4
On time (frames)
Norm. frequency
On-time (frames)
E
18 nm
F
Gray value
Distance (nm)
G
× 10⁴
Loc. Precision (nm)
Frequency
Photon yield
H
5 minutes
45 minutes

**Supplementary Fig. S3. 2D TDI-DNA-PAINT imaging of microtubules and DNA origami. (A)** The double-labeled ATTO Oxa14 probe (Ox2-R4) shows a ~6-fold increase upon addition of a $10^4$-fold excess of complementary docking strand 7×R4'. **(B)** Median background signal (photons/pixel/localization) of various concentrations of Ox1-P3 probe (green) were found to be 60 (100 pM), 122 (1 nM), 434 (5 nM), and 658 (10 nM). The same values for Ox2-P3 (orange) were 51.7 (100 pM), 55 (1 nM), 78 (5 nM), and 112 (10 nM). And for Ox2-R4 (magenta), the median background signal was 49 (100 pM), 54.5 (1 nM), 82 (5 nM), and 135 (10 nM) photons/pixel/localization. For Ox2-probes, we chose 5 nM as the optimal imager concentration for TDI-DNA-PAINT, and 100 pM for Ox1-P3 probes. **(C)** On-time distribution for Ox1-P3. Inset depicts a comparative plot of mean ± s.d. on-times (frames) for Ox1-P3 (1.8 ± 0.1), Ox2-P3 (1.9 ± 0.2), and Ox2-R4 (1.6 ± 0.2) as obtained from experiments on microtubule networks. **(D)** Comparative photon yields of Ox1-P3 (blue, 1,889 mean yield) and Ox2-P3 (orange, 3,545 mean yield) localizations. **(E)** DNA origami design. The distance between two adjacent docking sites is 18 nm. TDI-DNA-PAINT images show that 18 nm DNA origamis can be resolved after 5 min of imaging with 25 nM of Ox2-R4. Scale bars, 10 nm. **(F)** Line profile showing a peak-to-peak distance of 18.6 nm between two detected spots in the origami structure shown in (**E**). On average we detected a docking site distance of 18.9 ± 2.2 nm (SEM). **(G)** Photon yield distribution per binding event from Ox2-R4 (mean value: 4,183 photons) TDI-DNA-PAINT imaging of DNA origami. The inset shows the localization precision achieved of 3.9 ± 0.7 nm. (**H)** Typical TDI-DNA-PAINT images of 18 nm origami structures show gradually improved separation of docking sites over time due to accumulation of more localizations via imager re-binding kinetics. Scale bar, 10 nm.

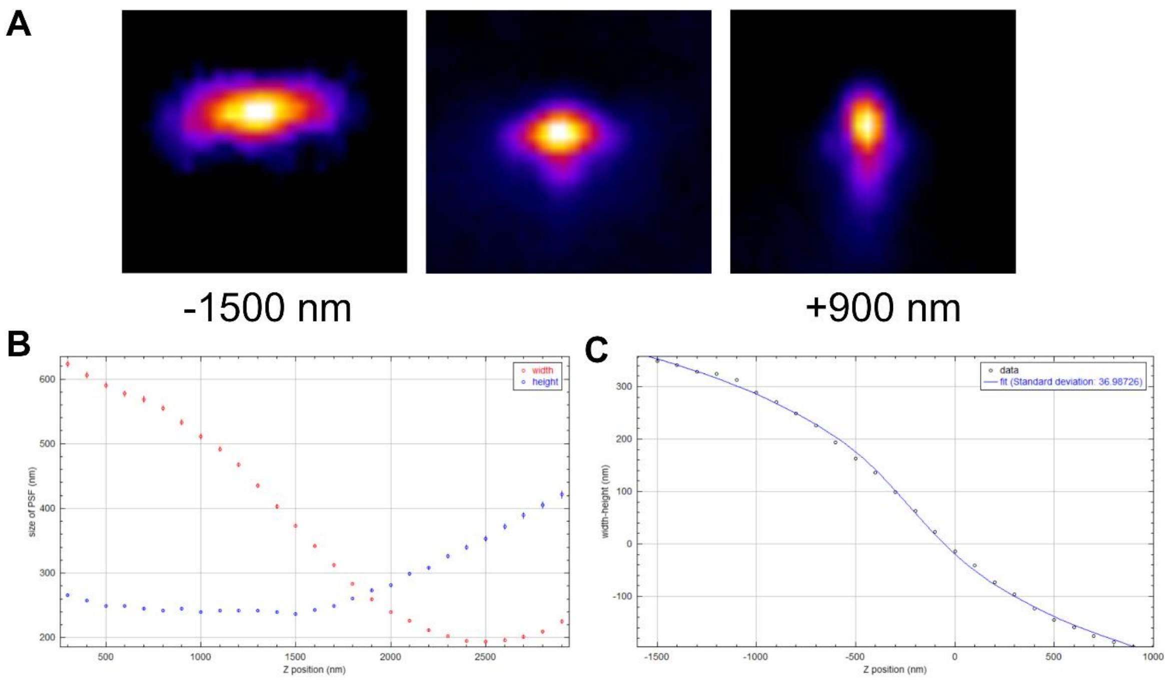


**Supplementary Fig. S4. LLS-TDI-DNA-PAINT PSF calibration**. **(A)** Average PSF (calculated from 5 individual beads) and its deformation along the z axis. Astigmatism and PSF shaping are achieved by proper tuning of the aberration correction parameter. **(B)** PSF size variation with z position. **(C)** Third degree polynomial fit.

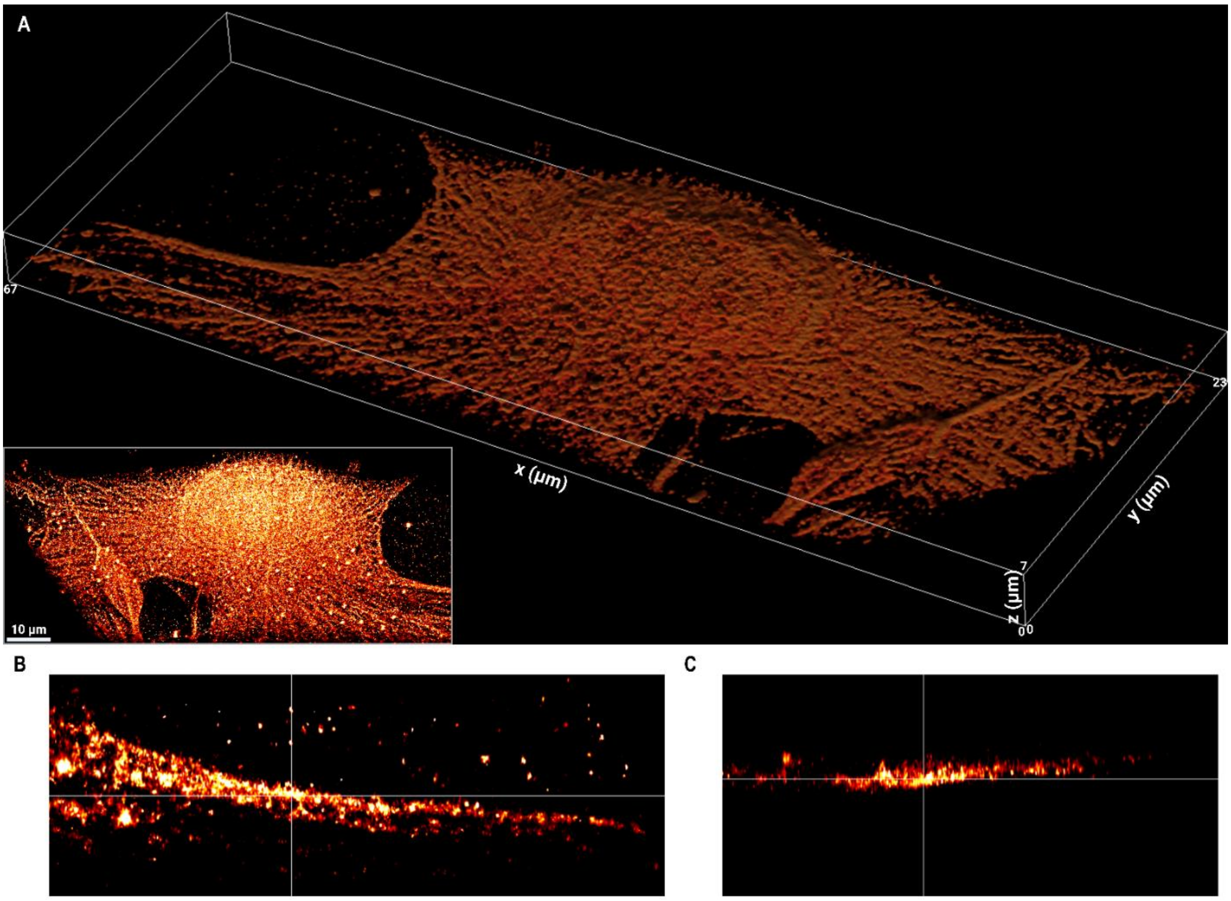


**Supplementary Fig. S5. LLS-TDI-DNA-PAINT imaging of COS7 cells. (A)** Whole-cell volume rendering (shadow projection) of a typical COS7 cell imaged with Ox2-P3 probe. The bounding box is 67 µm × 23 µm × 7 µm. Inset illustrates a maximum *z* projection of the same cell after deskewing but before rotation to the actual coverslip coordinates. (**B)** and **(C)** orthogonal views with *xy* (**B**) and *xz* **(C)** profiles of a representative ROI are shown.

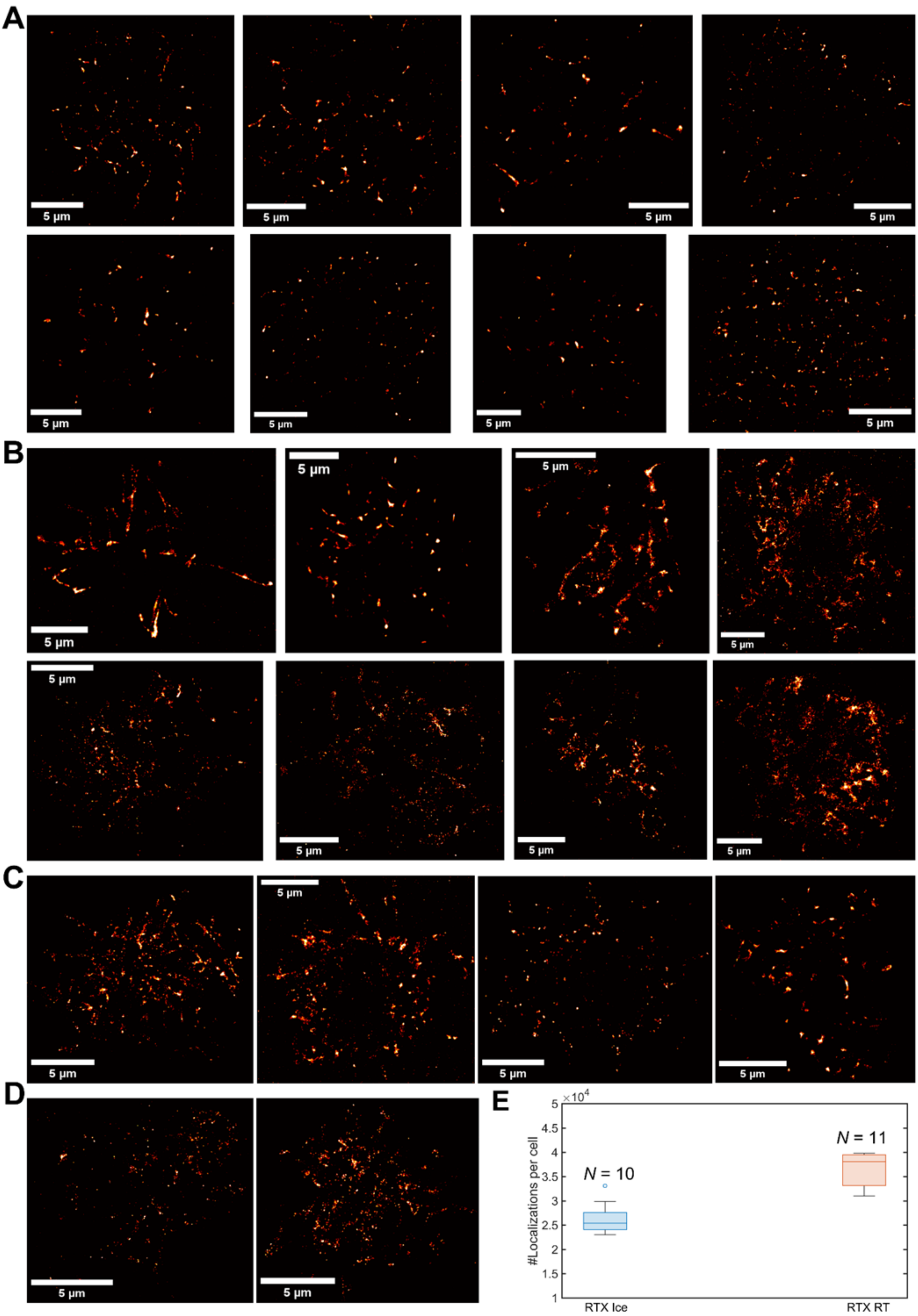


**Supplementary Fig. S6. 2D TDI-DNA-PAINT imaging of CD20 with RTX antibody in Raji B cells (A)** Nanoscale CD20 organization at the basal membrane of Raji cells. Cells were labeled with 7×R4' docking strand-tagged 5 µg/mL RTX. Labeling was performed on ice for 30 min. Distinct CD20 clusters localized in membrane protrusions/microvilli can be visualized. **(B)** Reconstructions for cells labeled with RTX and incubated for 30 min at RT exhibit prominent clusters and significantly longer (~1-8 microns) CD20-enriched protrusions as compared to (**A)**. **(C)** TDI-DNA-PAINT images of Raji cells with RTX incubated for 5 min and **(D)** 2 min at RT show CD20 clusters in microvilli of various dimensions. Longest spread-out protrusions and most densely compacted CD20 clusters are observed in case of **(B)** (30 min incubation at RT) confirming stabilization of membrane protrusions over time. **(E)** Quantification of number of localizations acquired in 10 min for cells labeled with RTX incubated at 4˚C and at RT. *N* denotes number of cells. Median number of detected localizations within 10 min were 38,100 and 25,400 for cells at RT and on ice, respectively.

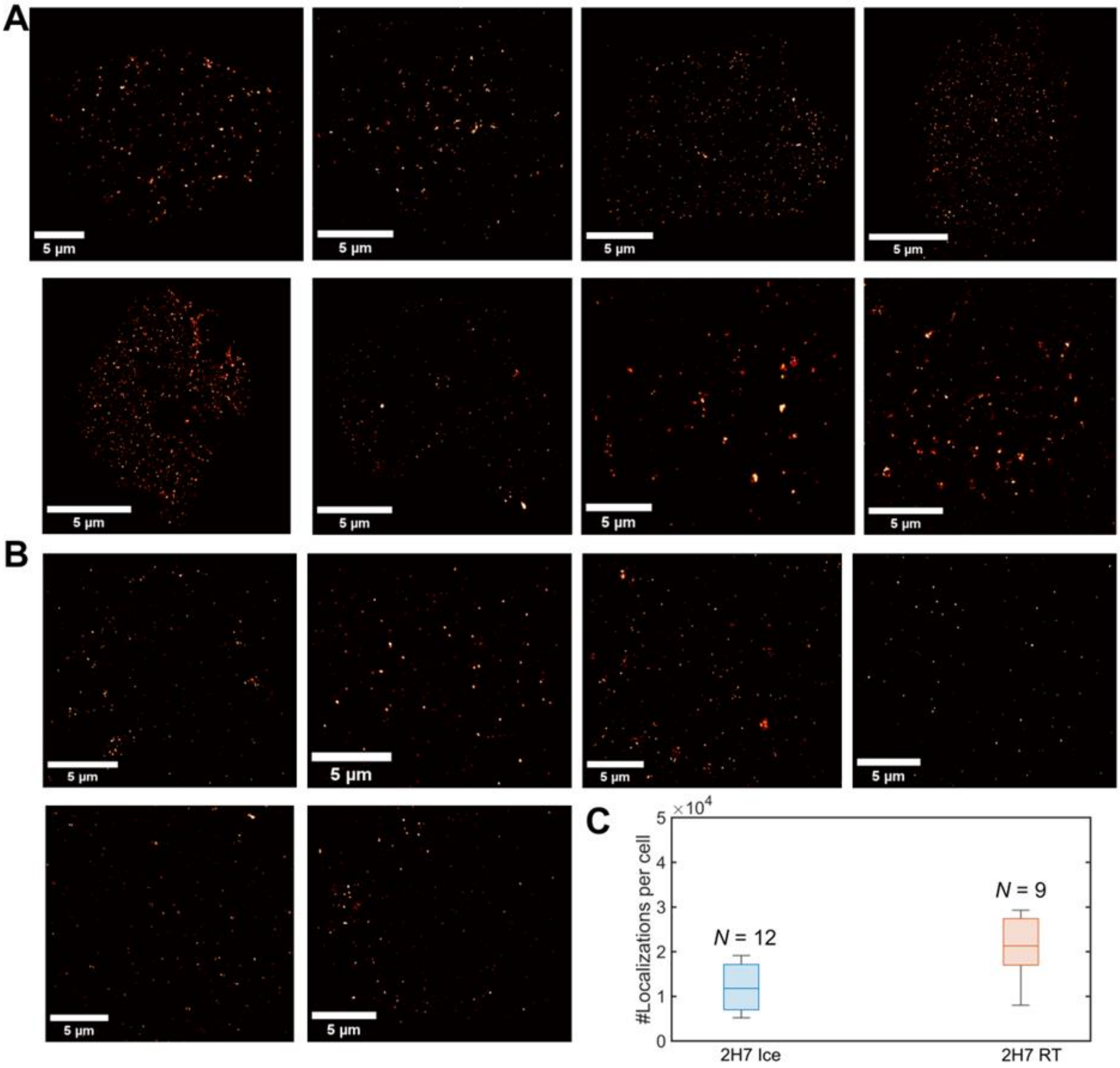


**Supplementary Fig. S7. 2D TDI-DNA-PAINT imaging of CD20 with 2H7 antibody in Raji B cells. (A)** CD20 organization at the basal membrane of Raji cells labeled with 5 μg/mL 2H7 and incubated for 30 min at RT. Oligomeric CD20 clusters are distributed across the basal membrane. **(B)** Reconstructions for cells labeled with 2H7 on ice incubated for 30 min (as shown in Fig. 3A in main text) exhibit lower abundance of CD20 clusters as compared to (**A**). **(C)** Quantification of number of localizations acquired in 10 min for cells labeled with 2H7 incubated at 4˚C (on ice) and at RT. *N* denotes number of cells. Median number of detected localizations were 21,300 and 11,800 for cells at RT and on ice respectively. Lower number of localizations relative to RTX-labeled cells (Fig. S6) can be explained by the lower binding affinity of 2H7.

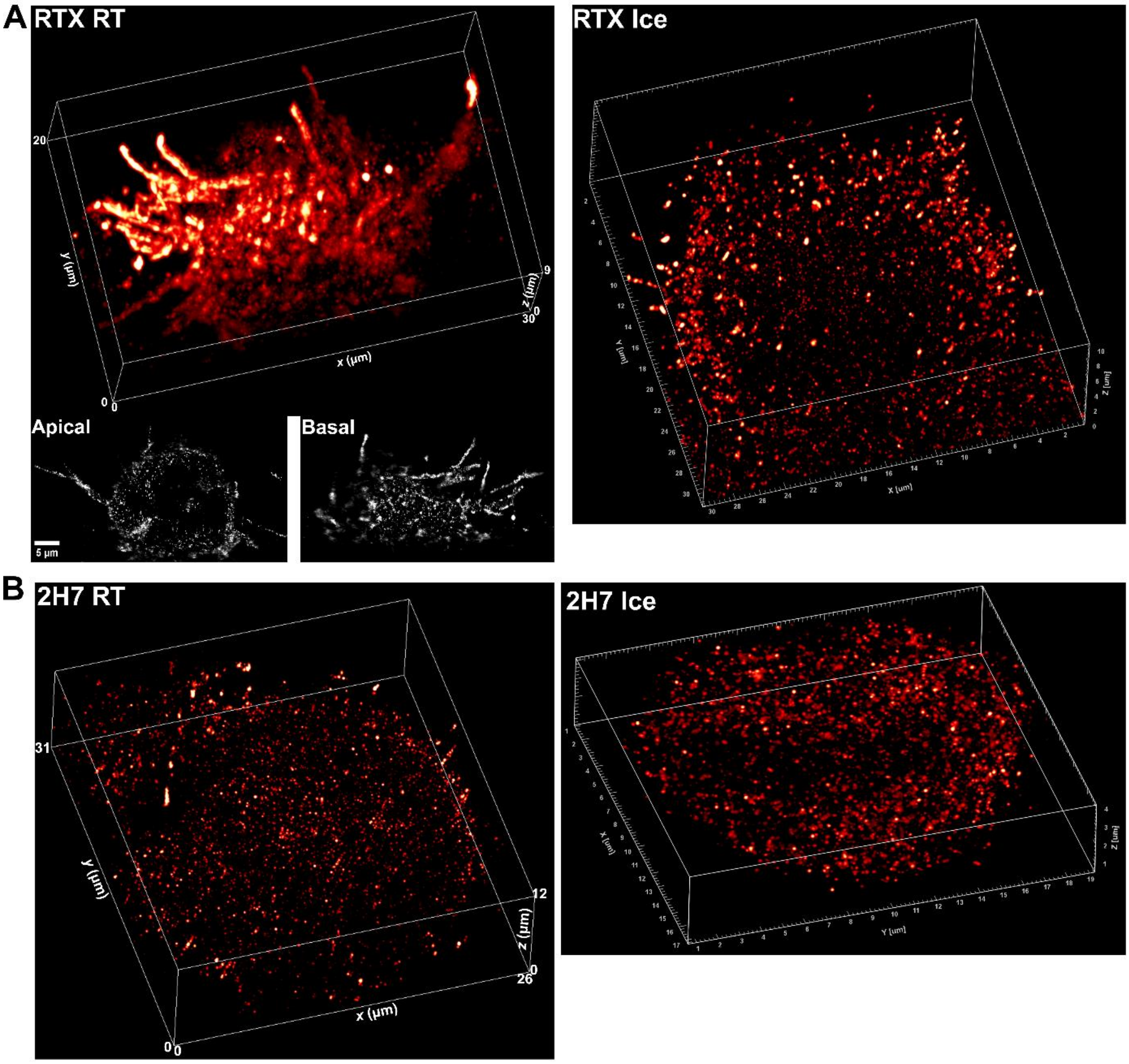


**Supplementary Fig. S8. LLS-TDI-DNA-PAINT imaging of CD20 in Raji cells. (A)** Whole-cell volumetric TDI-DNA-PAINT image of a typical Raji cell labeled for CD20 with 5 µg/mL RTX. Exemplary slices from apical and basal membranes are shown. RTX incubation was done for 30 min at RT and imaging was performed with Ox2-R4 probes (left). Clustering of CD20 into higher oligomeric states are visible, largely in membrane protrusions or microvilli spanning a few microns in length. Reconstructed Raji B cell volume showing RTX-CD20 distribution when incubation was done for 30 min on ice (right). **(B)** LLS-TDI-DNA-PAINT imaging of CD20 organization in Raji cell volumes where CD20 was labeled using 5 µg/mL anti-CD20 2H7 at RT (left) and for 30 min on ice (right).

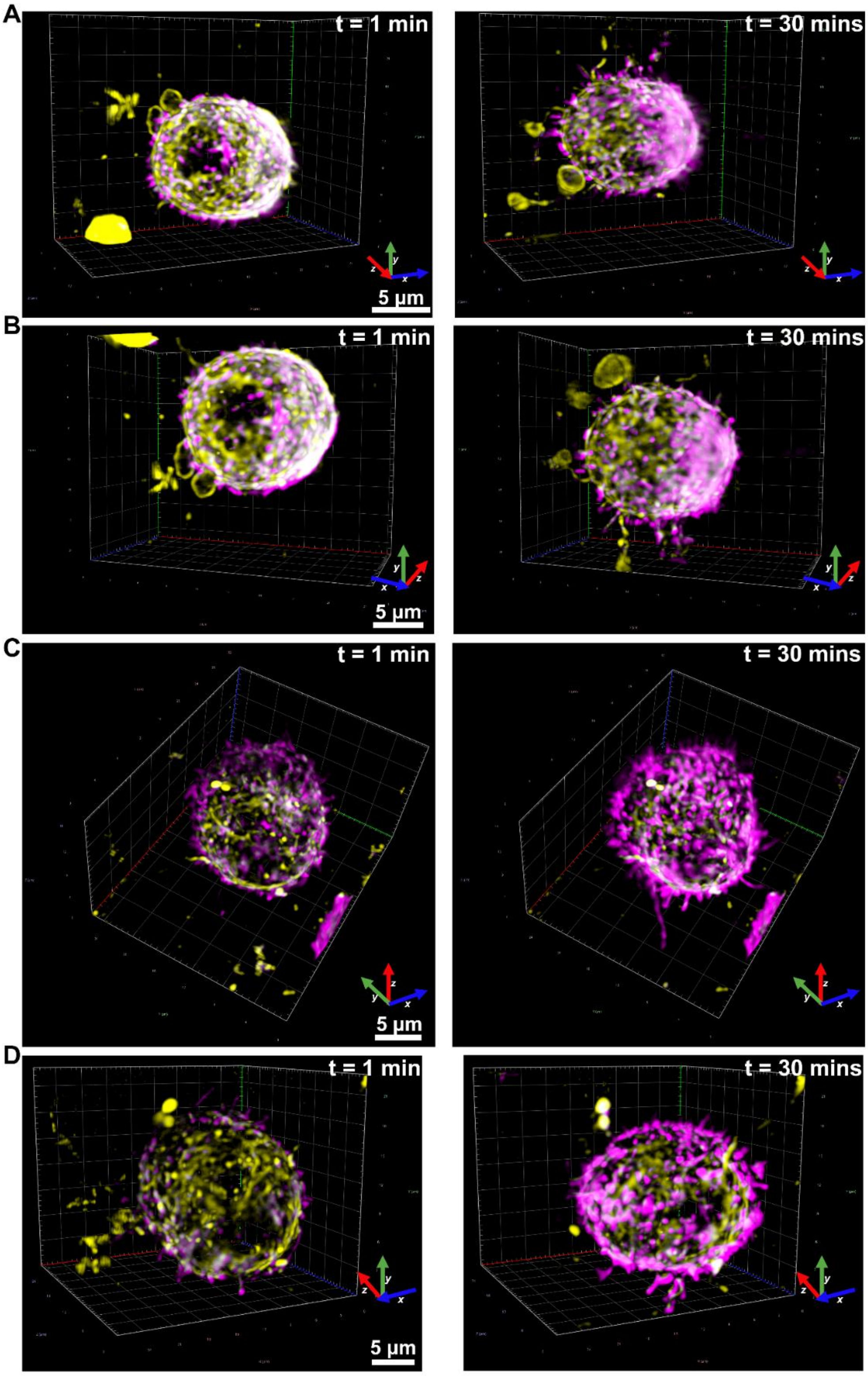


**Supplementary Fig. S9. Two-color live-cell LLS imaging of CD20 and actin in Raji B cells. (A)-(D)** Raji B cells were double stained with 5 µg/mL anti-CD20 RTX-AF647 (magenta) and live-cell dye SPY-555 for actin (yellow). Data acquisition was started right after addition of RTX to the sample chamber and cells were monitored for 30 min. Deconvolved LLS images at 1 and 30 min after antibody addition are shown. Cells manifest stabilization of microvilli of varied dimensions enriched in CD20 clusters along with polar CD20 aggregation and higher-order clusters. Displayed fluorescence signal gray value scale is same throughout for RTX and 2H7 channels, and for actin channel from figs. S9 to S13.

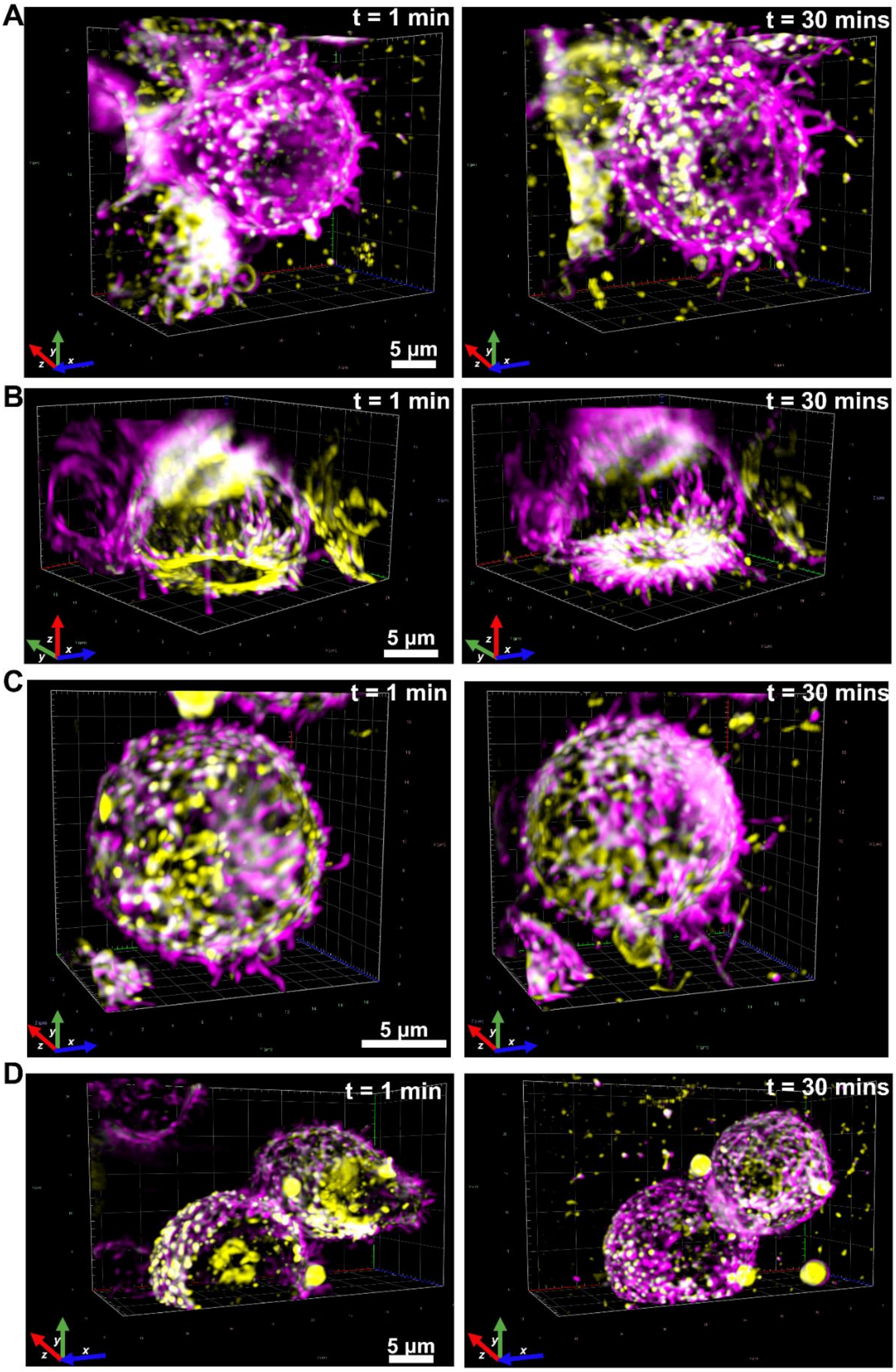


**Supplementary Fig. S10. Two-color live-cell LLS imaging of CD20 and actin in Raji B cells. (A)-(D)** Raji B cells were double stained with 10 µg/mL anti-CD20 RTX-AF647 (magenta) and live-cell dye SPY-555 for actin (yellow). Deconvolved LLS images at 1 and 30 min after antibody addition are shown. Same as in Fig. S9 cells manifest stabilization of microvilli of varied dimensions enriched in CD20 clusters along with polar CD20 aggregation and higher-order clusters. Displayed fluorescence signal gray value scale is same throughout for RTX and 2H7 channels, and for actin channel from figs. S9 to S13.

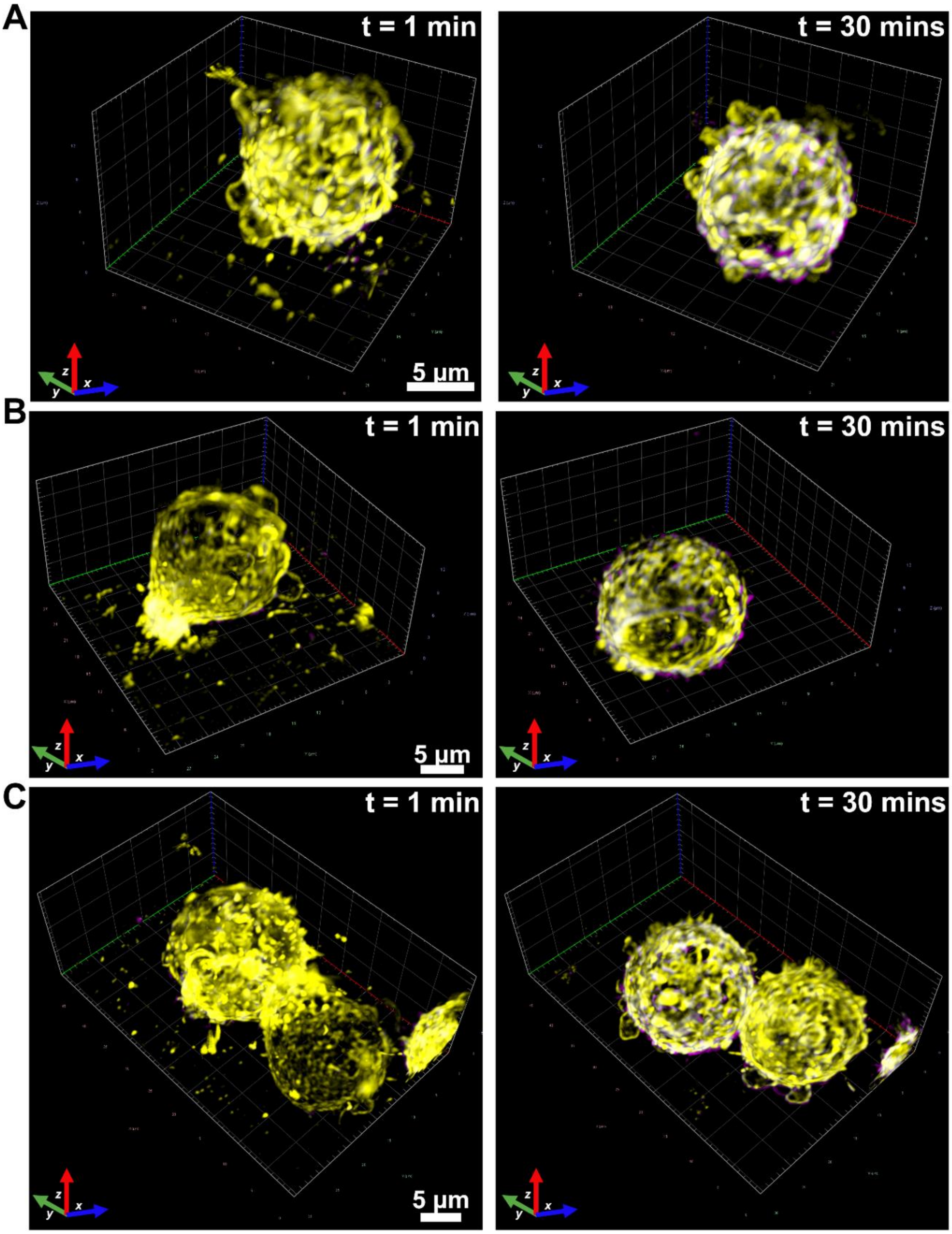


**Supplementary Fig. S11. Two-color live-cell LLS imaging of CD20 and actin in Raji B cells. (A)-(C)** Raji B cells were double stained with 5 μg/mL anti-CD20 2H7-AF647 (magenta) and live-cell dye SPY-555 for actin (yellow). Data acquisition was started right after addition of 2H7-AF647 to the sample chamber and cells were monitored for 30 min (same as the incubation time with RTX/2H7 for previously described TDI-DNA-PAINT experiments on Raji B cells). Deconvolved LLS images at 1 and 30 min after antibody addition are shown. Actin-rich membrane ruffles are visible. CD20 is distributed throughout the cell while the overall signal is significantly lower when compared to 5 μg/mL anti-CD20 RTX-AF647 (Fig. S9. Displayed fluorescence signal gray value scale is same throughout for RTX and 2H7 channels, and for actin channel from figs. S9 to S13.

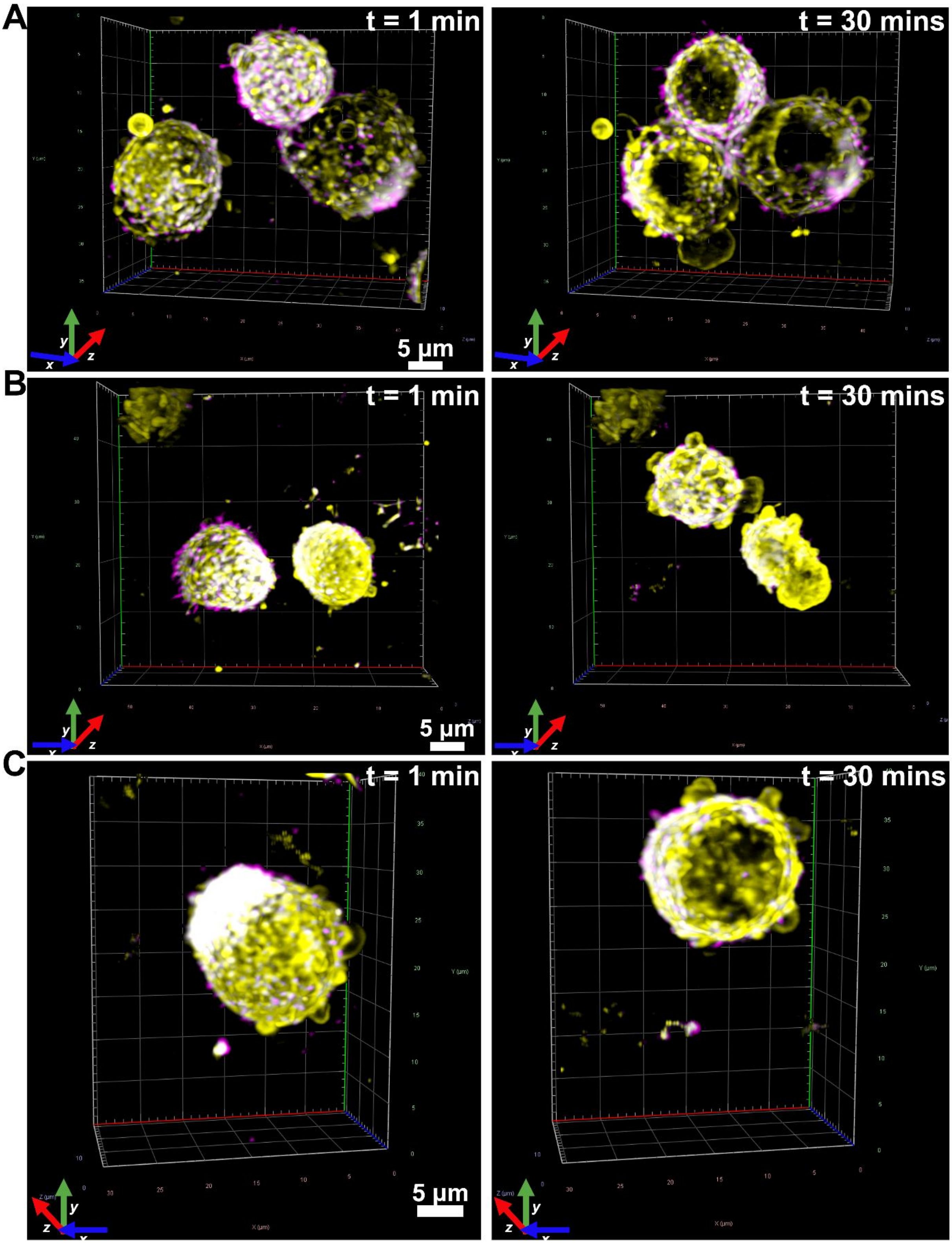


**Supplementary Fig. S12. Two-color live-cell LLS imaging of CD20 and actin in Raji B cells. (A)-(C)** Raji B cells were double stained with 10 µg/mL anti-CD20 2H7-AF647 (magenta) and live-cell dye SPY-555 for actin (yellow). Deconvolved LLS images at 1 and 30 min after antibody addition are shown. Actin-enriched membrane ruffles can be visualized. Cells show 'polarized' aggregation of CD20 and appearance of short membrane microvilli which are enriched with CD20. Displayed fluorescence intensity gray value scale is same throughout for RTX and 2H7 channels, and for actin channel from figs. S9 to S13.

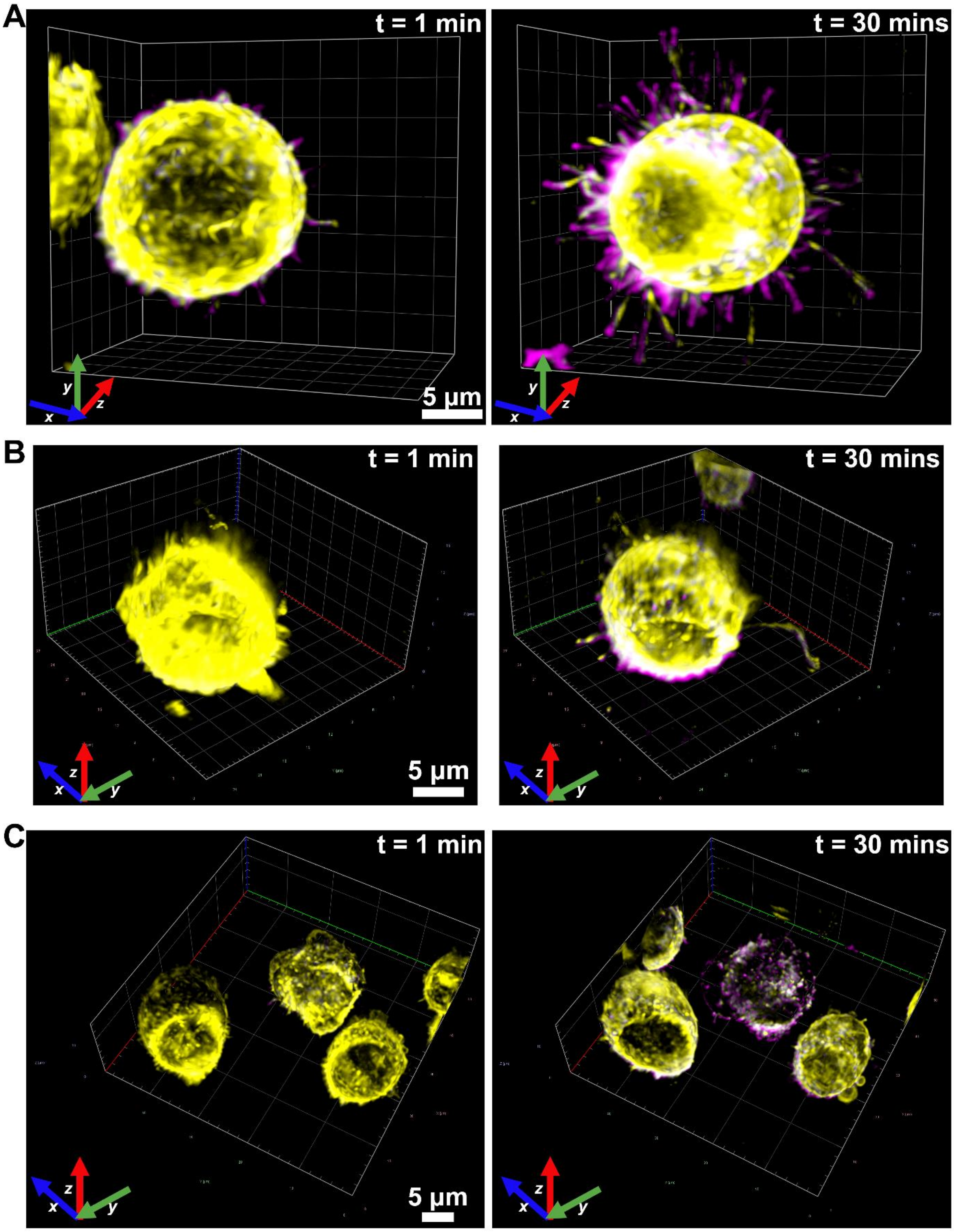


**Supplementary Fig. S13. Two-color live-cell LLS imaging of CD20 and actin in Raji B cells. (A)-(C)** Raji B cells were double stained with 20 µg/mL anti-CD20 2H7-AF647 (magenta) and live-cell dye SPY-555 for actin (yellow). Deconvolved LLS images at 1 and 30 min after antibody addition are shown. At t = 30 min, cells manifest 'polarized' accumulation and stabilization of membrane protrusions / microvilli and clustering of CD20. These observations indicate that at 20 µg/mL or even higher concentrations, 2H7 is also capable of inducing CD20 clustering and microvilli stabilization like RTX (5 µg/mL)-treated Raji cells (Figs.4A and S9. Displayed fluorescence signal gray value scale is same throughout for RTX and 2H7 channels, and for actin channel from figs. S9 to S13.

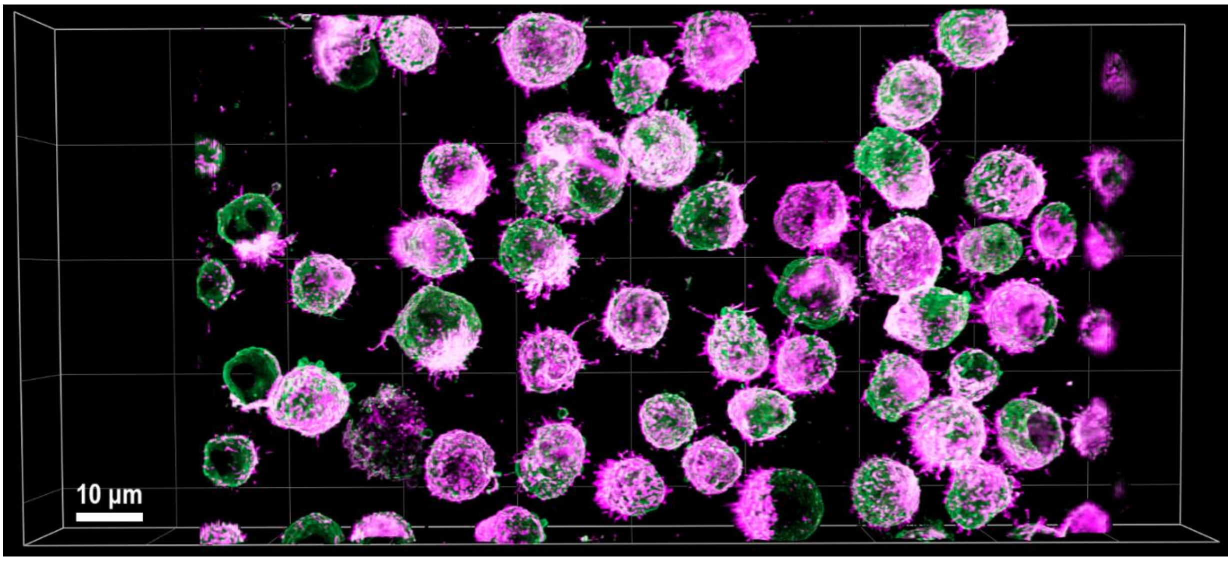


**Supplementary Fig. S14. Two-color LLS imaging of CD20 and CD45 in Raji B cells.** Raji B cells labeled with 5 µg/mL anti-CD20 RTX-AF647 (magenta) and anti-CD45 antibody-CF568 (green) incubated for 30 min at RT after RTX addition. Cells were fixed post-incubation. Deconvolved LLS images clearly demonstrate 'polarized' accumulation and clustering of CD20 in membrane protrusions, while CD45 is ubiquitously distributed throughout the cell membrane.

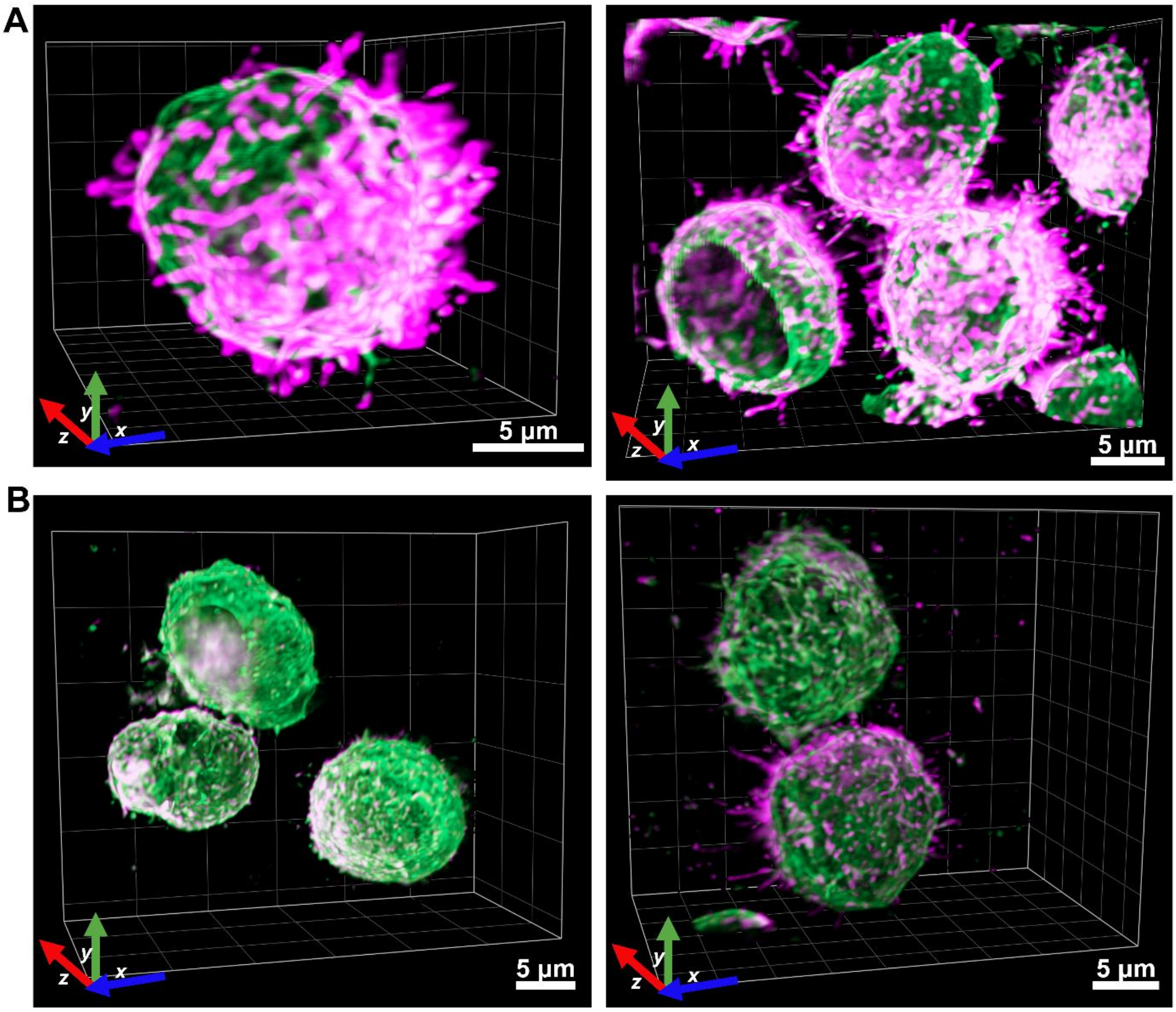


**Supplementary Fig. S15. Two-color LLS imaging of CD20 and CD45 in Raji B cells. (A)** Raji B cells labeled with 5 µg/mL anti-CD20 RTX-AF647 (magenta) and anti-CD45 antibody-CF568 (green) incubated for 30 min at RT after RTX addition. Cells were fixed post-incubation. Deconvolved LLS images demonstrate 'polarized' CD20 clusters and their accumulation in microvilli, while CD45 is distributed throughout the cell membrane. **(B)** Raji cells labeled with 5 µg/mL anti-CD20 2H7-AF647 (magenta) demonstrate significantly reduced signal globally as compared to **(A)** due to lower binding affinity of 2H7 and absence of strong CD20 clustering and stabilization of microvilli unlike RTX. Intensity gray value scaling was same for CD20-RTX and CD20-2H7 channels, and same for CD45 channels in **(A)** and **(B)**.

**Supplementary Movie S1:** Blinking movie of an exemplary 128 × 128 pixels area of DNA-PAINT imaging of microtubules using 100 pM Ox1-P3 imagers (Fig. 1D). Movie played at 50 fps.

**Supplementary Movie S2:** Blinking movie of an exemplary 128 × 128 pixels area of DNA-PAINT imaging of microtubules using 1 nM Ox1-P3 imagers. Movie played at 50 fps.

**Supplementary Movie S3:** Blinking movie of an exemplary 128 × 128 pixels area of DNA-PAINT imaging of microtubules using 5 nM Ox1-P3 imagers. Movie played at 50 fps.

**Supplementary Movie S4:** Blinking movie of an exemplary 128 × 128 pixels area of DNA-PAINT imaging of microtubules using 10 nM Ox1-P3 imagers. Movie played at 50 fps

**Supplementary Movie S5:** Blinking movie of an exemplary 128 × 128 pixels area of TDI-DNA-PAINT imaging of microtubules using 100 pM Ox2-P3 imagers. Movie played at 50 fps.

**Supplementary Movie S6:** Blinking movie of an exemplary 128 × 128 pixels area of TDI-DNA-PAINT imaging of microtubules using 1 nM Ox2-P3 imagers. Movie played at 50 fps.

**Supplementary Movie S7:** Blinking movie of an exemplary 128 × 128 pixels area of TDI-DNA-PAINT imaging of microtubules using 5 nM Ox2-P3 imagers (Fig. 1D). Movie played at 50 fps.

**Supplementary Movie S8:** Blinking movie of an exemplary 128 × 128 pixels area of TDI-DNA-PAINT imaging of microtubules using 10 nM Ox2-P3 imagers. Movie played at 50 fps.

**Supplementary Movie S9:** Blinking movie of an exemplary 128 × 128 pixels area of TDI-DNA-PAINT imaging of microtubules using 100 pM Ox2-R4 imagers. Movie played at 50 fps.

**Supplementary Movie S10:** Blinking movie of an exemplary 128 × 128 pixels area of TDI-DNA-PAINT imaging of microtubules using 1 nM Ox2-R4 imagers. Movie played at 50 fps.

**Supplementary Movie S11:** Blinking movie of an exemplary 128 × 128 pixels area of TDI-DNA-PAINT imaging of microtubules using 5 nM Ox2-R4 imagers (Fig. 1D). Movie played at 50 fps.

**Supplementary Movie S12:** Blinking movie of an exemplary 128 × 128 pixels area of TDI-DNA-PAINT imaging of microtubules using 10 nM Ox2-R4 imagers. Movie played at 50 fps.

**Supplementary Movie S13:** Blinking movie of an exemplary 128 × 128 pixels area of TDI-DNA-PAINT imaging of DNA origami using 25 nM Ox2-R4 imagers. Multiple origami structures (such as in figs. S2E and S2H) could be resolved in this area. Movie played at 50 fps.

**Supplementary Movie S14:** A fluorescent bead showing astigmatic aberration of the point spread function scanned over a range of -1500 to +900 nm at the LLS microscopy setup.

**Supplementary Movie S15:** Example blinking movie from LLS-TDI-DNA-PAINT imaging using Ox2-R4 (Fig. 2A) of a COS7 cell using 2.5 nM imager probes. Movie played at 25 fps.

**Supplementary Movie S16:** Deskewed z-stack of a COS7 cell after LLS-TDI-DNA-PAINT data analysis for Ox2-R4 imager. After deskewing, the cell is rotated to true coverslip coordinates as shown in Fig. 2A.

**Supplementary Movie S17:** Deskewed z-stack of a COS7 cell after LLS-TDI-DNA-PAINT data analysis for Ox2-P3 imager. After deskewing, the cell is rotated to true coverslip coordinates as shown in fig. S4.

**Supplementary Movie S18:** Blinking movie of an exemplary area of TDI-DNA-PAINT imaging of CD20 stained via 2H7 antibody on ice and imaged with 5 nM Ox2-R4 imagers (Fig. 3A). Movie played at 50 fps.

**Supplementary Movie S19:** Blinking movie of an exemplary area of TDI-DNA-PAINT imaging of CD20 stained via 2H7 antibody at RT and imaged with 5 nM Ox2-R4 imagers (Fig. 3A). Movie played at 50 fps.

**Supplementary Movie S20:** Blinking movie of an exemplary 256 × 256 pixels area of TDI-DNA-PAINT imaging of CD20 stained via RTX on ice and imaged with 5 nM Ox2-R4 imagers (Fig. 3A). Movie played at 50 fps.

**Supplementary Movie S21:** Blinking movie of an exemplary 256 × 256 pixels area of TDI-DNA-PAINT imaging of CD20 stained via RTX at RT and imaged with 5 nM Ox2-R4 imagers (fig. S5B). Movie played at 50 fps.

**Supplementary Movie S22:** Example blinking movie from LLS TDI-DNA-PAINT imaging in a Raji cell where CD20 was labeled with RTX and imaged with 5 nM Ox2-R4 imagers. Movie played at 50 fps.

**Supplementary Movie S23:** Example blinking movie from LLS TDI-DNA-PAINT imaging in a Raji cell where CD20 was labeled with 2H7 and imaged with 5 nM Ox2-R4 imagers. Movie played at 50 fps.

**Supplementary Movie S24:** Reconstructed LLS-TDI-DNA-PAINT image stack of a Raji cell labeled with RTX (Fig. 3B).

**Supplementary Movie S25:** Reconstructed LLS-TDI-DNA-PAINT image stack of a Raji cell labeled with anti-CD20 2H7 (Fig. 3C and fig. S7B).

**Supplementary Movie S26:** Two-color LLS imaging on Raji cells showing 5 µg/mL RTX-stained CD20 (magenta) and actin (yellow) dynamics for 30 min after the start of experiment.

**Supplementary Movie S27:** Two-color LLS imaging on Raji cells showing 5 µg/mL 2H7-stained CD20 (magenta) and actin (yellow) dynamics for 30 min after the start of experiment.

**Supplementary Movie S28:** Two-color LLS imaging on Raji cells showing 10 µg/mL RTX-stained CD20 (magenta) and actin (yellow) dynamics for 30 min after the start of experiment.

**Supplementary Movie S29:** Two-color LLS imaging on Raji cells showing 10 µg/mL 2H7-stained CD20 (magenta) and actin (yellow) dynamics for 30 min after the start of experiment.

**Supplementary Movie S30:** Two-color LLS imaging on Raji cells showing 20 µg/mL 2H7-stained CD20 (magenta) and actin (yellow) dynamics for 30 min after the start of experiment.
